# A redox switch in p21–CDK feedback during G2 phase controls the proliferation-cell cycle exit decision

**DOI:** 10.1101/2024.09.14.613007

**Authors:** Julia Vorhauser, Theodoros I. Roumeliotis, David Coupe, Jacky K. Leung, Lu Yu, Kristin Böhlig, Thomas Zerjatke, Ingmar Glauche, André Nadler, Jyoti S. Choudhary, Jörg Mansfeld

## Abstract

Reactive oxygen species (ROS) influence cell proliferation and fate decisions by oxidizing cysteine residues (S-sulfenylation) of proteins, but specific targets and underlying regulatory mechanisms remain poorly defined. Here, we employ redox proteomics to identify cell cycle-coordinated S-sulfenylation events and investigate their functional role in proliferation control. Although ROS levels rise during cell cycle progression, overall oxidation of the proteome remains constant with dynamic S-sulfenylation restricted to a subset of cysteines. Among these, we identify a critical redox-sensitive cysteine residue (C41) in the cyclin-dependent kinase (CDK) inhibitor p21. C41 oxidation regulates the interaction of p21 with CDK2 and CDK4, controlling a double-negative feedback loop that determines p21 stability. When C41 remains reduced, p21’s half-life increases in G2 phase resulting in more p21 inheritance to daughter cells suppressing proliferation and promoting senescence after irradiation. Notably, we identify dynamic S-sulfenylation on further cell cycle regulators implying coordination of cell cycle and redox control.

## INTRODUCTION

Reactive oxygen species (ROS) have long been known for their ability to damage proteins, lipids, and DNA. At physiological levels however, ROS, especially membrane-permeant hydrogen peroxide (H_2_O_2_) function in a concentration-dependent manner as signaling molecules in processes such as proliferation, differentiation, and stress responses^1^. H_2_O_2_ is mainly derived from superoxide anions produced by the mitochondrial electron transport chain and NADPH oxidases and converted to H_2_O_2_ by superoxide dismutases. The main cellular targets of ROS are proteins with sulfur-containing amino acids cysteine and methionine due to their high oxidation rate constants compared to lipids and nucleic acids^2^. Because H_2_O_2_ oxidizes cysteine ∼4x faster than methionine, reversible cysteine oxidation plays a predominant role in redox signaling^3^.

Upon reaction with H_2_O_2_, cysteine forms a sulfenic acid (S-sulfenylation, SOH), which can further react to form disulfides or conjugates with thiol-containing metabolites such as glutathione (glutathionylation) or acetyl coenzyme A (CoAthiolation or CoAlation)^4^. These modifications are reversible via thioredoxin or glutathione-dependent systems, emphasizing their regulatory role in protein function, localization, stability, and interactions^5^.

Evidence increasingly links reversible cysteine oxidation (oxPTMs) to cell proliferation. ROS levels rise during the cell cycle^6–9^, and blocking ROS production or enhancing antioxidant activity can delay G1- or S phase^6,9^, or inhibit mitosis^10,11^. Proteomic approaches using differential alkylation or sulfenic acid probes have identified hundreds of oxPTMs in core cell cycle regulators^12–14^, yet the timing and functional impact of these modifications remain poorly understood.

Recently, cell cycle kinases have identified as key targets of ROS: Oxidation of Aurora A regulates its activation^4,15,16^, CDK4 forms redox-sensitive disulfide-linked interactions with cyclin D that reduce proliferation^17^; and oxidation of CDK2 near its T-loop modulates its regulation by phosphatases and kinases^9^. Whether redox control operates in a coordinated, cell cycle–dependent manner across the proteome remains an open question.

CDK activity drives cell cycle progression and is controlled by T-loop phosphorylation and interactions with activators or inhibitors, respectively. The timely activation of CDKs during G1 phase determines whether cells commit to proliferation or exit the cell cycle into quiescence^18^. The CDK inhibitor p21 (CDKN1A), which inhibits CDK4/6-cyclin D, CDK2-cyclin A/E, and CDK1-cyclin B1 complexes, plays a critical role in this decision-making^19–21^. Cells entering mitosis with high levels of p21, e.g., due to DNA damage or replication stress, give rise to daughter cells destined for cell cycle exit. Therefore, p21 levels are tightly controlled before mitosis by a double-negative feedback loop involving CDK2 and the SCF-SKP2 E3 ubiquitin ligase^21,22^. CDK2-mediated phosphorylation of p21 creates a phospho-degron that promotes recognition by SCF-SKP2, leading to proteasomal degradation of p21^23–25^. Conversely, elevated p21 levels inhibit CDK2 activity, thereby preventing its own phosphorylation and further stabilizing p21. To prevent unphysiologically high concentrations of p21 during G2 phase, which affect cell fate of daughter cells, it is critical to maintain dynamic negative feedback between p21 and CDK2. How this balance is achieved, and the molecular mechanism(s) involved, remain unknown.

Here, we performed sulfenic acid–specific proteomics at 2-hour intervals across the cell cycle and identified 1725 unique S-sulfenylation sites in non-transformed retinal pigment epithelial (RPE-1) cells. Although global S-sulfenylation remains stable, oxidation of specific cysteines including cysteine 41 (C41) of p21 was dynamic. C41 oxidation peaked in G2 phase, and we show that the redox state of C41 at this time regulates p21 stability. The reduced form of C41 strengthens p21’s inhibition of CDK2 in G2 phase, stabilizing p21 and shifting the p21-CDK2 feedback in favour of p21. Under high p21 expression (e.g., following radiation), this redox switch biases cells toward cell cycle exit and senescence.

## RESULTS

### Cell Cycle-Resolved Redox Proteomics Identify Dynamic S-Sulfenylation

To label sulfenylated cysteines in living RPE-1 cells, we used the recently developed, cell-permeant probe BTD (Figure S1A)^12,26^. BTD irreversibly labels sulfenic acids of living cells with high reactivity (1,700 M⁻¹ s⁻¹) and contains an alkyne moiety for click chemistry (Figure S1B). Live-cell labelling with BTD for one hour followed by conjugation to an azide-fluorophore (Az488) using click chemistry revealed strong sulfenic acid signals in all compartments, including nucleus and ER/mitochondria-like structures (Figure S1C), confirming widespread sulfenic acid formation in RPE-1 cells.

Previous work, including our own, has shown that ROS levels rise during the cell cycle^6,7,9^. To test whether sulfenic acids similarly increase, we performed BTD labelling followed by flow cytometry and observed elevated sulfenylation in S and G2/M phases compared to G1 (Figure S1D), consistent with earlier findings^7,27^. To determine whether these changes reflect proteome-wide S-sulfenylation or specific oxidation events, we developed a quantitative redox-proteomic workflow. RPE-1 cells were synchronized by release from 150 nM palbociclib^28^ and sulfenic acids were labelled with BTD at multiple time points for one hour. Proteins were extracted and biotinylated via azide-picolyl-Dde-biotin using click chemistry. Following digestion, peptides were enriched on streptavidin beads, labelled with tandem mass tags (TMT)^29^ and eluted by Dde cleavage prior to mass spectrometry (MS). Parallel total proteome samples were used to normalize S-sulfenylation to total protein abundance (Figure 1A). Flow cytometry confirmed high synchronicity between biological repeats (Figure 1B), supported by consistent changes in known cell cycle regulators CDT1, cyclin A2, and geminin (Figure S1E). To focus on physiologically relevant stages, early time points (0 h, 3 h), which may be affected by palbociclib treatment, were excluded in favor of later unperturbed G1 time points (17 h, 19 h). Across all cell cycle stages, we identified 1,725 unique S-sulfenylation sites, the highest number reported for BTD in living, unperturbed cells (Table S1). Despite an increase in S-sulfenylation in flow cytometry, normalization to protein abundance revealed no overall increase during the cell cycle (Figure 1C). To identify functionally relevant changes, we filtered for sites whose oxidation profiles matched characteristic S, G2/M, or G1 patterns, yielding 233 dynamic cysteines (Figure 1D; Table S1). Gene ontology analysis showed G2/M-high sites were enriched in cytoskeletal and cell cycle proteins, including CFL, IQGAP1, CLASP2, PCM1, KIF23, and p21 (highlighted in black). Most dynamic sites mapped to cytosolic proteins, consistent with proximity to ROS-producing organelles (Figure 1E; Table S1).

**Figure 1.**
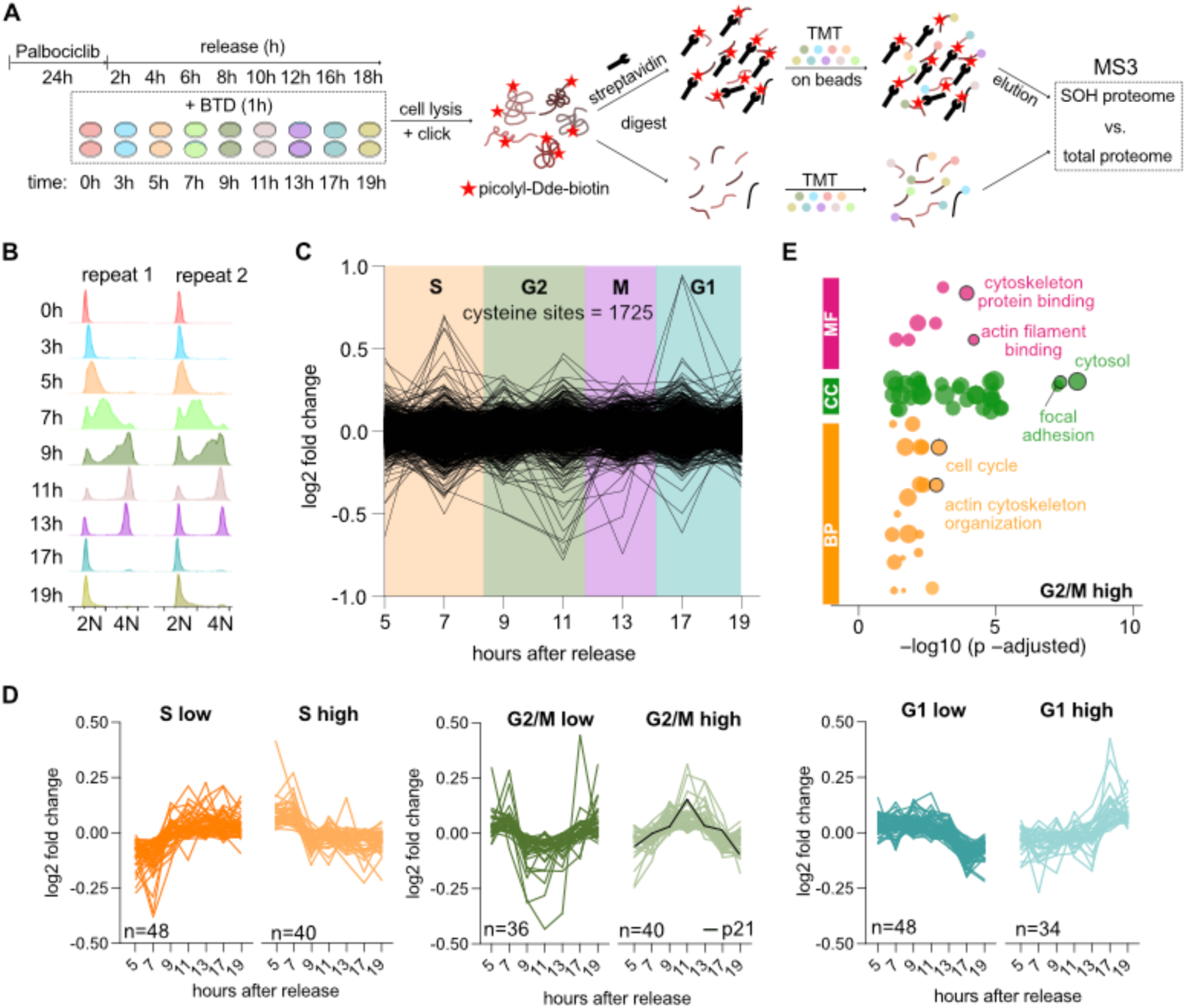
Cell Cycle-Resolved Redox Proteomics Identify Dynamic S-Sulfenylation. (**A**) Proteomic workflow to identify cell cycle-dependent S-sulfenylation using BTD in RPE-1 cells. (**B**) DNA content of BTD-treated RPE-1 cells prior to mass spectrometry, determined by flow cytometry. (**C**) Cell cycle-dependent changes in S-sulfenylation normalized to protein abundance; each line represents a unique oxidized cysteine. (**D**) Clustering of dynamic S-Sulfenylation sites into prototypical cell cycle oscillations (see Table S1). p21 S-Sulfenylation is highlighted in black (G2/M high). (**E**) GO enrichment analysis for Biological Process (BP), Molecular Function (MF), and Cellular Component (CC) of proteins in the G2/M-high cluster. GO analyses for other clusters and full annotation are provided in Table S1.

Together, these data show that while overall proteome sulfenylation remains constant across the cell cycle, a subset of cysteines undergo dynamic, site-specific S-sulfenylation suggesting regulatory roles analogous to phosphorylation.

### p21 is Oxidized at C41 During G2 Phase

Among core cell cycle regulators, the CDK inhibitor p21 (CIP/KIP family) exhibited dynamic S-sulfenylation across the cell cycle. To confirm p21 oxidation, asynchronous RPE-1 cells were treated with BTD for 2 hours, followed by lysis and click reaction to biotin-azide and enrichment on streptavidin beads (Figure 2A). p21 and the known redox-regulated protein GAPDH^30^ were enriched on BTD-treated but not DMSO-treated beads. Based on input-to-elution signal ratios, p21 appeared more oxidized than GAPDH at steady state. As expected, RPL26, which lacks cysteines, was absent on BTD-treated beads.

**Figure 2.**
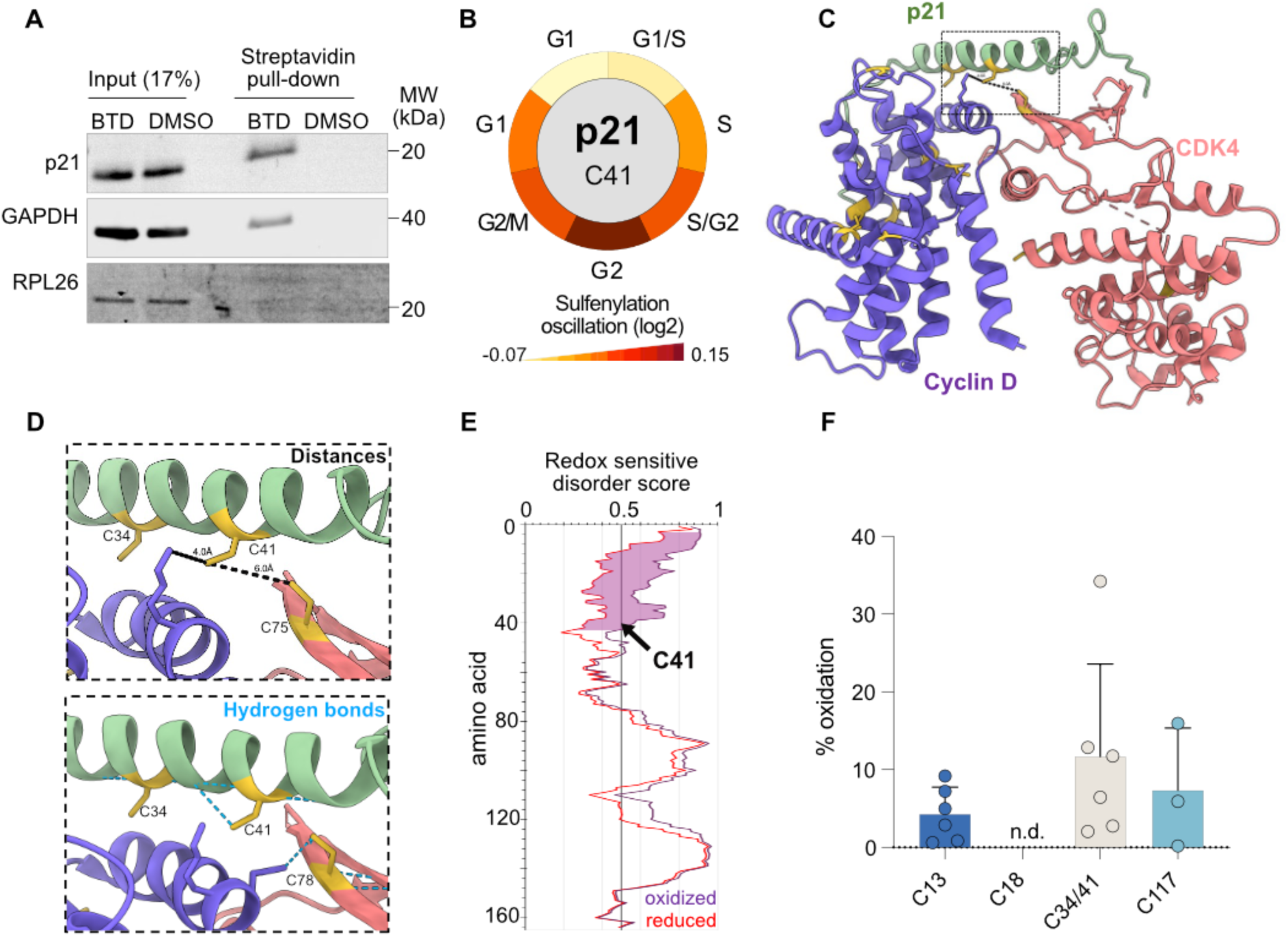
p21 is Oxidized on C41 During G2 Phase. (**A**) Western blot from asynchronous RPE-1 cells showing streptavidin pull-downs of S-sulfenylated proteins (p21, GAPDH) labelled with BTD and subsequently clicked to azide-biotin, (n=2). (**B**) Cell cycle-dependent S-Sulfenylation of p21 identified by BTD proteomics (see also Table S1). (**C**) Structure of the p21 (green)-CDK4 (red)-cyclin D (purple) complex (PDB: 6p8h^32^). (**D**) Enlargement of the molecular interface around C41 from panel (C) showing distance measurements (top) and hydrogen bonds (bottom). (**E**) Prediction of order-to-disorder transitions upon p21 oxidation using IUPred3. (**F**) Stoichiometry of p21-GFP oxidation in G2 phase, determined by MS (see workflow in Figure S2E). C34 and C41 are located within the same peptide and cannot be resolved individually. Bars represent mean ± SD, n=6.

Cell cycle-resolved BTD proteomics revealed S-sulfenylation of p21 at C41, peaking in G2 phase (Figures 1D, 2B). C41 is highly conserved in vertebrates and unique among CIP/KIP proteins (Figures S2A, S2B). Though p21 is intrinsically disordered, its N-terminal domain (aa 9–85) adopts an α-helical conformation upon CDK–cyclin binding^31^. Structural analysis of p21–CDK4–cyclin D^32^ and a homology model of p21–CDK2–cyclin A^33^ placed C41 at the interface of all three proteins (Figures 2C, S2C). C41 lies close to cyclin D (4 Å), CDK4 (6 Å), and cyclin A (2.7 Å), and forms a hydrogen bond with the α-helix backbone, suggesting a role in helix stabilization (Figures 2D, S2C)^34^. Notably, p21 adopts a more extended conformation when bound to CDK2–cyclin A, altering the position of C41 and potentially affecting complex specificity (Figure S2D)^35^. IUPred3 predictions indicate the N-terminal region, including C41, undergoes order-to-disorder transitions upon oxidation, consistent with redox-sensitive structural plasticity (Figure 2E)^36^.

While BTD redox proteomics showed that C41 oxidation peaks in G2 phase, this approach only provides relative, not absolute (stoichiometry), information about C41’s oxidation state. To quantify the stoichiometry of C41 oxidation in G2, we used differential alkylation-reduction-alkylation MS combined with p21 enrichment (Figure S2E)^14,37^. RPE-1 cells expressing endogenous p21-GFP^21^ were synchronized in G2 by an 11-hour palbociclib release (Figure 1B). Following selective cysteine labelling and GFP pulldown^38,39^ MS analysis identified oxidation at four of five cysteines of p21: C13 (4%), C117 (7%), and C34/C41 (12%) (Figure 2F). While C34 and C41 reside on the same peptide and thus could not be resolved, BTD proteomics exclusively detected oxidation at C41 (Figure 2B, Table S1), suggesting this site is the primary target of oxidation.

We conclude that p21 is predominantly oxidized at C41, with oxidation peaking in G2 phase.

### Mutating C41 Increases p21 Protein Levels and Impairs Proliferation

To assess whether C41 oxidation affects p21 function in living cells, we mutated C41 to serine (C41S) to mimic the reduced, non-oxidizable state. *In vitro* kinase assays with recombinant CDK2–cyclin A and its substrate Retinoblastoma protein (RB) (aa 773–928), supplemented with recombinant p21-WT, C41S, or C41A, revealed similar IC_50_’s under reducing conditions (Figure S3A). This confirmed that mutagenesis did not impair p21 inhibitory function. Given its structural similarity to cysteine (Figure S3B), we selected C41S for cell-based studies.

Using CRISPR–Cas9 with a co-selection strategy^40^ we generated two homozygous C41S knock-in clones in RPE-1 cells expressing endogenous p21-GFP^21^, confirmed by amplicon-next generation sequencing (NGS) (Figure S3B-C). Fluorescence imaging under unperturbed conditions showed a mild but significant increase in p21-positive cells in C41S mutants (WT: 7%; C41S #1: 11%; C41S #2: 10%) (Figure 3A–B), with slightly higher p21-GFP intensities (Figure S3D). γH2AX staining and Western blotting confirmed no increase in DNA damage or p53 expression (Figure 3C, S3E–F). Similar findings were observed in CRISPR-edited mammary breast epithelial cells (MCF10A) cells, where p21 levels were strongly elevated in heterozygous and homozygous C41S clones, while p53 remained unchanged (Figure S3G–J). The stronger increase in p21 levels upon C41S mutagenesis suggests that MCF10A cells are more sensitive to p21 perturbations than RPE-1 cells^21,41^. Consequently, we were unable to maintain stable p21-C41S lines in MCF10A cells.

**Figure 3.**
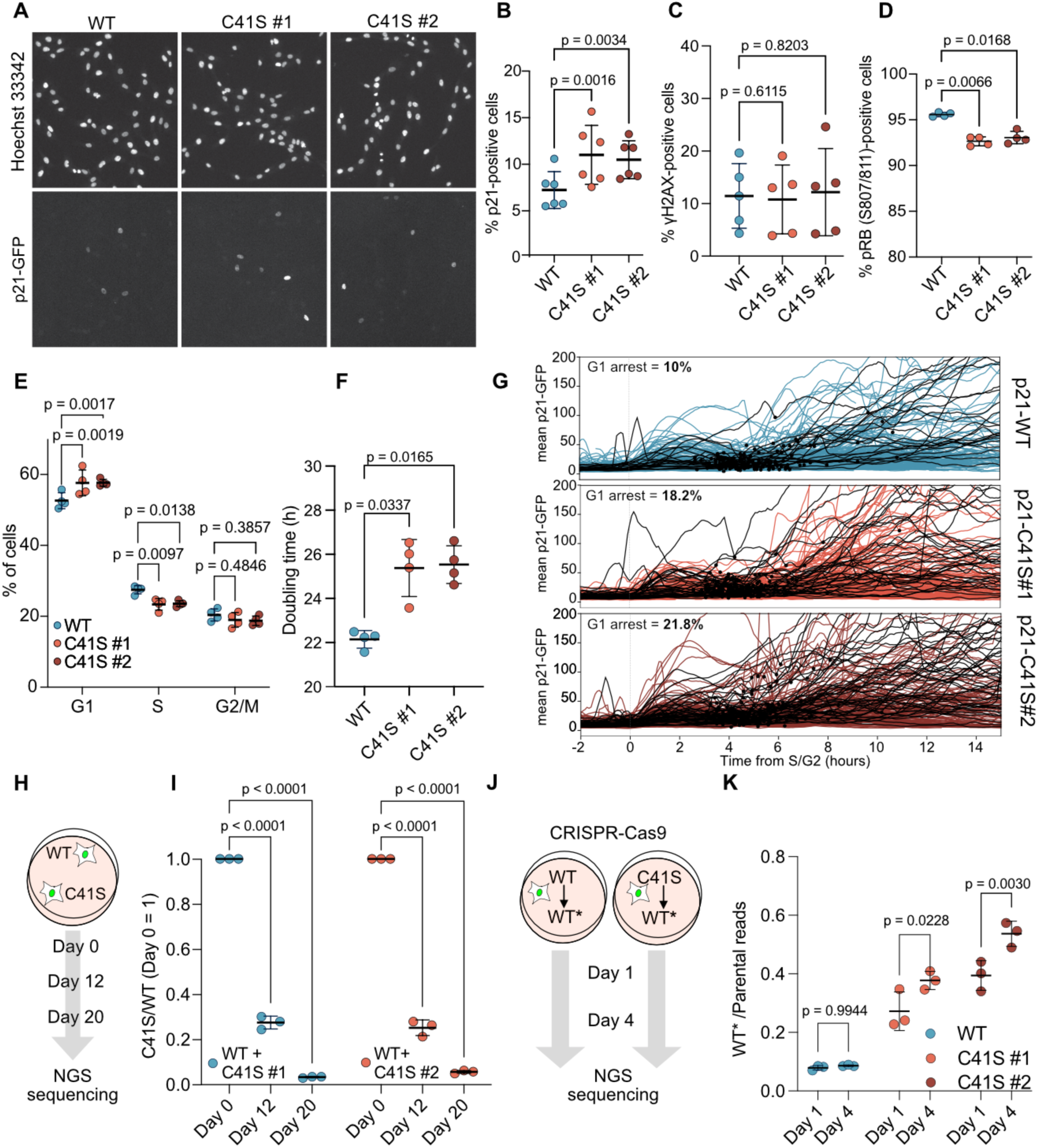
Mutating C41 Increases p21 Protein Levels and Impairs Proliferation. (**A**) Imaging of p21-GFP expression and DNA (Hoechst 33342) in RPE-1 cells showing parental (WT) and two independent C41S clones. Scale bar, 50 µm. (**B**) Quantification of p21-positive cells from experiments shown in (A); mean ± SD, n = 6. Significance by one-way ANOVA. (**C**) Quantification of γH2AX-positive p21-GFP WT and C41S cells (mean ± SD; n = 5). Representative images shown in Figure S3E. Significance by one-way ANOVA. (**D**) Quantification of pRB S807/811-positive cells (mean ± SD; n=4). Significance by one-way ANOVA. (**E**) Cell cycle distribution of p21-GFP WT and C41S cells based on Hoechst 33342 staining (mean ± SD; n = 4). Significance by two-way ANOVA. (**F**) Doubling time of p2-GFP WT and C41S cells (mean ± SD; n = 2, N = 4). Significance by one-way ANOVA. (**G**) Single-cell tracks from time-lapse imaging of p21-GFP, C41S #1, and C41S #2 cells aligned at S/G2 phase transition. Black dots indicate mitoses and black lines mark cell tracks with G1 phases >15 h (n = 2, N = 220). (**H**) Schematic of competitive proliferation assay between p21-GFP WT and C41S cells. (**I**) Ratio of C41S/WT reads at days 0, 12, and 20 of co-culture as outlined in (H) (mean ± SD; n = 3). Significance by two-way ANOVA. (**J**) Schematic of CRISPR-Cas9-mediated reversion of C41S or WT (C41S→WT*) and WT to WT (WT→WT*) using a NGS-detectable repair template (WT*). (**K**) Ratio of WT*/parental reads at days 1 and 4 post-reversion (mean ± SD; n = 3). Significance by two-way ANOVA.

Since p21 inhibits CDK-mediated RB phosphorylation (pRB)^42–44^, we measured pRB S807/811 levels. While Western blotting showed no obvious differences (Figure S3F), more sensitive immunofluorescence revealed a significant reduction in pRB-positive cells in C41S clones (Figure 3D). Cell cycle profiling showed G1-phase accumulation and S-phase reduction in C41S cells (Figure 3E, S3K), alongside a ∼3.5 h longer cell cycle duration (Figure 3F). Single-cell time-lapse imaging of asynchronously growing p21-GFP cells, classified by mRuby-PCNA^45^, showed that C41S mutants exhibit higher p21 levels during G2 and mitosis, leading to greater inheritance of p21 in G1 daughter cells (Figure 3G, S3L–N). The fraction of daughter cells with extended G1 (>15 h) doubled (WT: 10%; C41S #1: 18.2%; C41S #2: 21.8%), suggesting that preventing C41 oxidation extents G1 phase or causes cell cycle exit.

To evaluate the long-term effects of the inability to oxidize C41, we co-cultured WT and C41S cells and tracked genotype frequencies by amplicon-NGS. Already after 12 days, C41S cells contributed only 25% of reads and were nearly undetectable by day 20 (<3%) (Figure 3H-I). To exclude clonal variation or CRISPR-Cas9 off-target effects as cause for the loss of proliferative capacity in C41S cells, we reverted C41S to WT (C41S>WT*) and introduced synonymous mutations in WT cells (WT>WT*) as a control. Amplicon-NGS tracking showed a significant increase in C41S>WT* cells over four days (C41S #1: 27%→38%; C41S #2: 40%→55%), while WT>WT* remained stable (∼10%) (Figure 3J–K). The increase of C41S>WT* compared to WT>WT* cells on day one post-electroporation likely reflects that C41S cells, which already exhibit elevated steady-state p21 levels, are hypersensitive to electroporation-induced p53 activation and cell death^46^.

In summary, preventing C41 oxidation increases p21 levels in G2, resulting in elevated inheritance to daughter cells and increased G1 duration/cell cycle exit. Although these effects are subtle short-term, they become strongly disadvantageous under long-term or competitive conditions, underscoring the importance of C41 redox regulation for p21 homeostasis and sustained proliferation.

### C41 Redox State Regulates p21 Stability by Increased CDK2 Inhibition

After observing that cells with non-oxidizable p21 C41S exhibit increased p21 levels (Figure 3B) and impaired proliferation (Figure 3I), we investigated the molecular mechanism underlying this effect. C41 is centrally positioned the p21-CDK-cyclin-interface (Figure 2D, S2C), suggesting its oxidation could influence p21’s interaction with CDK-cyclin complexes. To test this, we examined whether p21 WT and C41S differed in their ability to bind CDK-cyclin complexes. We prepared lysates from asynchronously growing RPE-1 WT and C41S cells and performed p21-GFP pull-down assays using GFP-binder beads, omitting reducing reagents during lysis and pull-down to preserve potential C41 oxidation. Western blotting for bound CDK-cyclin complexes revealed that p21-GFP C41S bound significantly less CDK4-cyclin D1 compared to p21-GFP WT but showed increased binding to CDK2 (Figure S4A-D). Interactions of p21 with cyclin A2, cyclin B1, and CDK1 were comparable between WT and C41S cells (Figure S4E-G). We noticed that, unlike CDK2, cyclin A2 was not significantly enriched on p21-GFP C41S possibly reflecting that in asynchronous cells CDK2 also binds to cyclin E and cyclin A to CDK1 potentially decreasing the sensitivity of this assay. Importantly, adding the reducing reagent DTT during pull-down assays abrogated the differences in CDK4 and CDK2 binding between p21-GFP WT and C41S, in agreement with their equal inhibition of CDK2 activity *in vitro* under reducing conditions (Figure S4H-J, S3A). This suggests that the redox state of C41, rather than the C41S mutation itself, alters p21-CDK-cyclin interactions.

Because in asynchronous cells G2 phase is underrepresented, we next synchronized cells in G2 phase, where C41 oxidation peaks (Figure 2B), and confirmed less CDK4 but more CDK2 binding by p21-C41S (Figure 4A–C). Treating cells with 10 µM H₂O₂ for 30 minutes before lysis increased CDK4 binding to p21-GFP WT but not C41S, without affecting p21-CDK2 binding.

**Figure 4.**
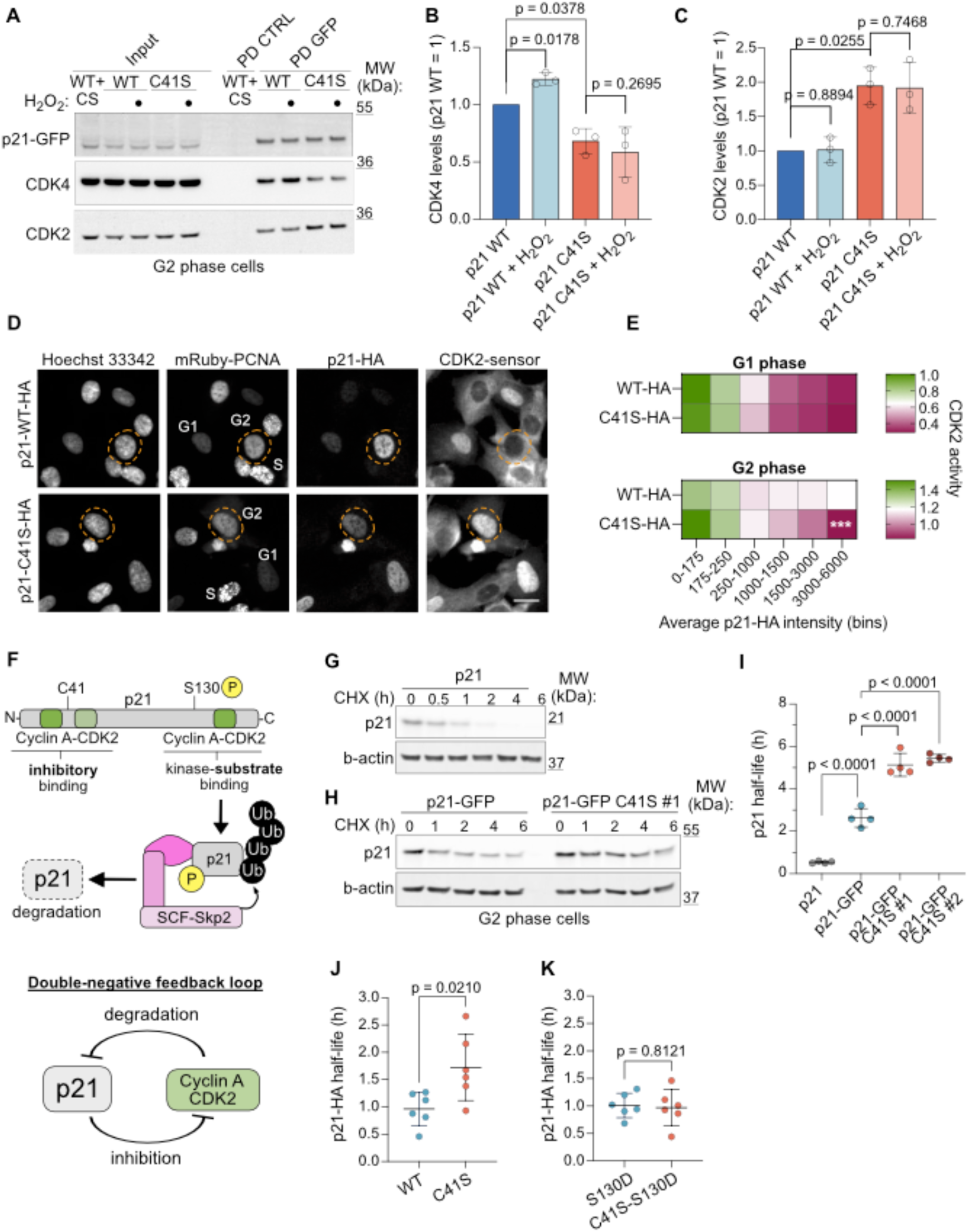
p21 Stability and Inheritance Depends on C41 Oxidation. (**A**) Western blot of control (CTRL) and GFP pull-downs from RPE-1 p21-GFP cells synchronized in G2, detecting CDK2 and CDK4 interactions in the presence or absence of 10µM H_2_O_2_. (**B, C**) Quantification of CDK4 (B) and CDK2 (C) bound to p21-GFP WT or C41S#1 based on data in (A) (mean ± SD, n=3). Significance by one-sample (p21 WT vs. p21 WT+H_2_O_2_, p21 WT vs. p21 C41S) and paired t-tests (p21 C41S vs. p21 C41S+H_2_O_2_). (**D**) Immunofluorescence of RPE-1 cells expressing p21-WT-HA or p21-C41S-HA, endogenously tagged mRuby-PCNA, and a CDK2 activity sensor that localizes to the cytoplasm when CDK2 is active. G2 cells were identified by PCNA localization and are outlined (dashed lines). Scale bar, 20 μm. (**E**) Quantification showing mean CDK2 activity per bin during G1 and G2 in RPE-1 WT and C41S cells grouped by p21 expression, based on data from (D). Cell cycle stage was determined by DNA content (Hoechst 33342). Significance by to two-way ANOVA. (***p = 0.0002; n = 4). (**F**) Schematic of p21 interaction with CDK2–cyclin A and its phosphorylation-dependent recognition by the SCF–SKP2 ubiquitin ligase, indicating inhibitory and substrate interactions and the resulting double-negative feedback loop. (**G**) Western blot of RPE-1 cells synchronized in G2 and treated with cycloheximide (CHX) for the indicated times to assess p21 half-life. β-actin serves as loading control. (**H**) Western blot as in (G) comparing half-lives of endogenous p21-GFP WT and C41S#1. (**I**) Quantification of p21, p21-GFP, and p21-GFP C41S (clones #1 and #2) half-lives, based on decay curves shown in Figure S4L. Statistical significance by one-way ANOVA (mean ± SD; n = 3, N = 4). (**J, K**) Quantification of half-lives of p21-WT-HA and p21-C41S-HA (J), and phospho-mimetic mutants p21-WT-S130D-HA and p21-C41S-S130D-HA (K) during G2, based on decay curves shown in Figures S4N and S4O. Statistical significance by one-way ANOVA (mean ± SD; n = 6).

To test whether stronger CDK2 binding by p21-C41S enhances CDK2 inhibition, we used a CDK2 activity sensor in a p21⁻/⁻ RPE-1 background expressing mRuby-PCNA and tetracycline-inducible p21-human influenza hemagglutinin (HA)^47^. Stable lines expressing p21-WT-HA or p21-C41S-HA at physiological levels (Figure S4K) revealed that CDK2 activity was lower in C41S-expressing G2 cells, as indicated by nuclear retention of the sensor (Figure 4D). Using anti-HA immunofluorescence and DNA content-based gating, we binned G1 and G2 cells by p21 levels and quantified CDK2 activity. While the degree of CDK2 inhibition correlated with p21 levels in G1 for both WT and C41S cells, C41S cells showed significantly stronger CDK2 inhibition in G2 at equivalent p21 levels (Figure 4E). Consistent with p21-GFP pull-downs this suggests that p21 with reduced C41 is the better CDK2 inhibitor (Figure 4A-C).

p21 interacts with CDK2 both as an inhibitor and as a substrate whereby CDK2 phosphorylates p21 on S130, promoting SCF-SKP2–mediated degradation (Figure 4F)^23–25^. This establishes a double-negative feedback loop, suggesting that enhanced CDK2 inhibition by C41S may impair p21 degradation. To test this, we blocked protein synthesis with cycloheximide in G2-arrested cells and monitored p21 decay by Western blot. The half-life of untagged p21 was ∼30 min^48^, while p21-GFP extended this to ∼2.5 h (Figure 4G–I, S4L). The C41S mutation further stabilized p21-GFP half-life to >5 h. This effect was specific to G2-phase cells because no difference was observed in asynchronous populations (Figure S4M), where alternative degradation pathways dominate^21,49,50^. Accordingly, p21-C41S inhibited CDK2 more effectively than p21-WT in G2 but not in G1 phase (Figure 4D, E).

To confirm that this stabilization was due to impaired S130 phosphorylation, we introduced a phospho-mimetic S130D mutation into p21-WT-HA and p21-C41S-HA and expressed both to endogenous levels in RPE-1 p21⁻/⁻ cells (Figure S4K). In G2-phase cells, treated with CHX to block translation, p21-C41S-HA showed a half-life of ∼1.7 h, nearly double that of p21-WT-HA (∼1 h), while the C41S+S130D mutant restored degradation to WT kinetics (Figure 4J–K, S4N–O).

We conclude that the redox state of C41 regulates p21’s interaction with CDK2 and CDK4. When C41 cannot be oxidized, the double-negative feedback between p21 and CDK2 shifts in favor of p21, reducing CDK2 activity, and slowing p21 degradation. C41 oxidation likely functions upstream of S130 phosphorylation, linking redox regulation to CDK activity and p21 stability.

### C41 Redox State Sensitizes Cells to Stress-Induced p21 Accumulation

The prolonged half-life of p21-C41S in G2 phase (Figure 4I) suggested enhanced sensitivity to stress-induced p21 expression. To test this, we treated p21-GFP WT and C41S cells with various p21-inducing stimuli and quantified single-cell p21-GFP levels in G2 cells (identified by Hoechst 33342 DNA content). As expected, in unperturbed conditions p21 peaked in G1 and was low in S, with variable levels in G2/M (Figure S5A)^21^. After low-dose ionizing radiation (2 Gy) and 24 h recovery, C41S cells showed a markedly larger p21^high^ G2 population than WT (Figures 5A–B). Similar effects were observed following Nutlin-3 treatment to induce p53 expression (1 µM, 24 h), which also increased the proportion of G2 cells with low pRB S807/811 staining from 5% in WT to 17% and 14% in C41S mutants (Figures 5C–E, S5B), consistent with reduced CDK2 activity in G2 (Figure 4E).

**Figure 5.**
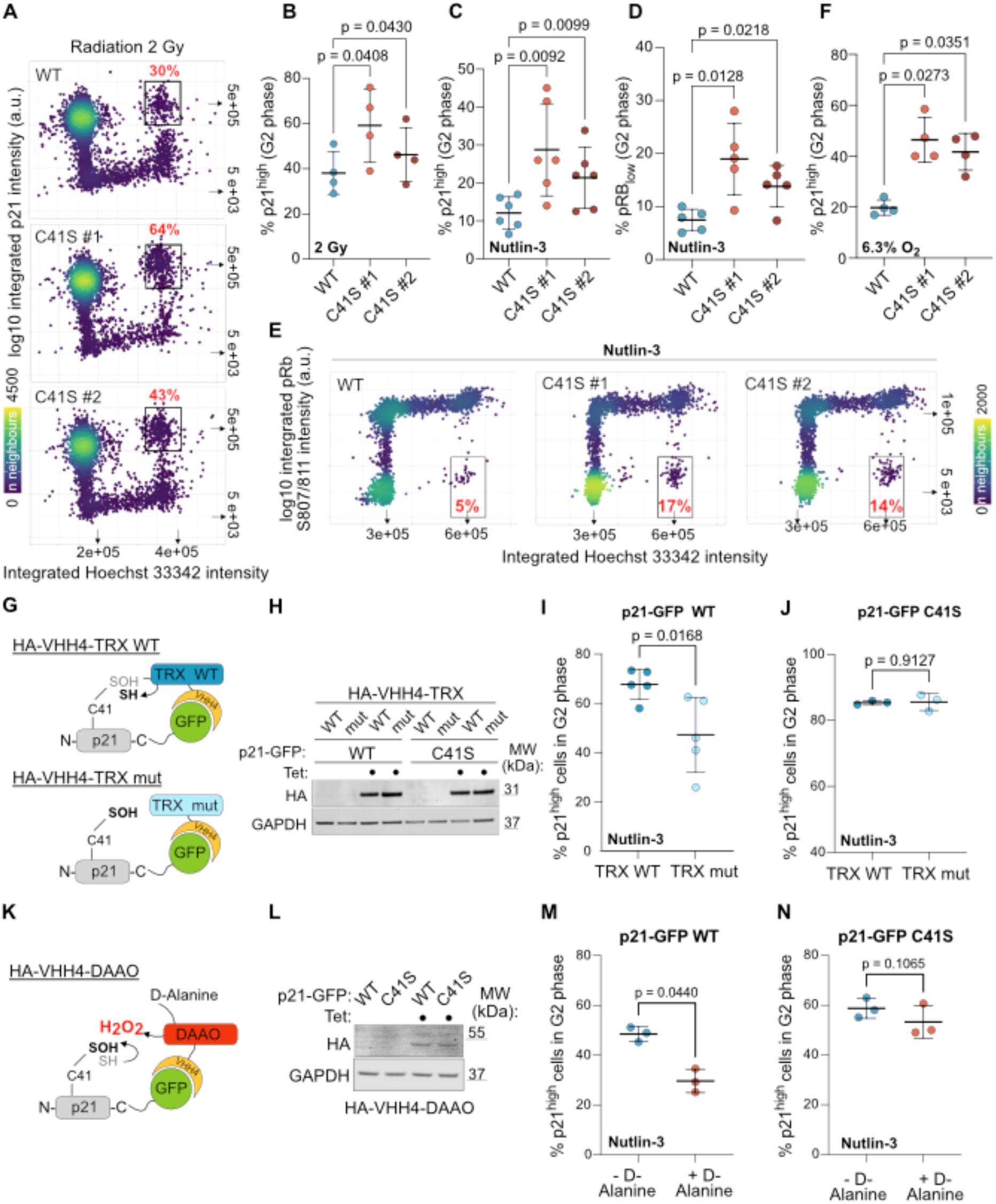
C41 Redox State Modulates p21 Stability and Sensitivity to Cellular Stress. (**A**) Bivariate density plots showing p21 levels and DNA content (Hoechst 33342) in RPE-1 p21-GFP WT and C41S cells 24 h after irradiation with 2 Gy. Gates and percentages for p21^high^ cells in G2 are indicated. Corresponding untreated controls are shown in Figure S5A. **B**) Quantification of p21^high^ cells in G2 from (A). Significance by one-way ANOVA (mean ± SD; n = 4). (**C**) Percentage of p21-GFP p21^high^ cells in G2 after treatment with 1 µM Nutlin-3 for 24 h. Significance by one-way ANOVA (mean ± SD; n = 6). (**D**) Quantification of pRB S807/811 staining from (E). Significance by one-way ANOVA (mean ± SD; n = 5). (**E**) Bivariate density plots showing phospho-RB (pRB S807/811) staining in RPE-1 p21-GFP WT and C41S cells after 24 h of 1 µM Nutlin-3 treatment. Gates and percentages for G2 cells with low pRB staining are indicated. Corresponding untreated cells are shown in Figure S5B. (**F**) Percentage of p21-GFP p21^high^ cells in G2 after 9–11 days of culture at 6.3% O₂, a condition that induces DNA damage and p21 expression (see Figure S5C–F). Significance by one-way ANOVA (mean ± SD; n = 3, N = 4). (**G**) Strategy to maintain p21 in a reduced state using tetracycline-inducible expression of a GFP nanobody (VHH4) fused to either wild-type (WT) or catalytically inactive (mut) thioredoxin (TRX), targeting antioxidant activity to p21-GFP. (**H** Western blot showing comparable expression of HA-VHH4-TRX WT and HA-VHH4-TRX mut in RPE-1 p21-GFP cells as used for (I, J). GAPDH serves as loading control. (**I, J**) Percentage of p21^high^ cells in G2 in p21-GFP WT (I) and p21-C41S-GFP (J) cells expressing HA-VHH4-TRX WT or mut after 24 h of 1 µM Nutlin-3 treatment. Statistical significance by paired t-test (mean ± SD; n = 5 for I, n = 3 for J). Corresponding bivariate density plots and gating are shown in Figures S5G and S5H. (**K**) Strategy to maintain p21 in an oxidized state using tetracycline-inducible expression of GFP nanobody VHH4 fused to D-amino acid oxidase (DAAO), producing H₂O₂ near p21-GFP in response to D-alanine addition. (**L**) Western blot showing comparable expression levels of HA-VHH4-DAAO in RPE-1 p21-GFP cells as used for (M, N). GAPDH serves as loading control. (**M, N**) Percentage of p21^high^ cells in G2 in p21-GFP WT (M) and p21-C41S-GFP (N) cells expressing HA-VHH4-DAAO after 3 h of 5 µM Nutlin-3 treatment. Statistical significance by paired t-test (mean ± SD; n = 3). Bivariate plots used for gating are shown in Figures S5J and S5K.

Since RPE-1 cells cultured at 21% O₂, experience hyperoxia relative to their native environment, we also assessed p21 regulation under physiological oxygen (6.3% O₂)^51,52^. After ≥5 days of adaptation, p21-positive cells increased from <10% (21% O₂), to >60% (6.3% O₂) (Figures S5C–D), coinciding with increased γH2AX staining (Figure S5E–F). Under these conditions, C41S mutants again displayed nearly double the G2 p21^high^ population compared to WT (Figure 5F).

To further corroborate that our observations are due to the redox state of C41 rather than the C41S mutation *per se*, we developed a system to specifically modulate the redox state of p21 in living RPE-1 cells. We fused a GFP nanobody (VHH4)^38,39^ to the antioxidant enzyme Thioredoxin (TRX) to target its reducing activity directly to p21-GFP and prevent p21 oxidation. To this end we generated isogenic tetracycline-inducible stable cell lines in the p21-GFP WT and C41S background expressing wild-type (TRX WT) or a catalytically inactive mutant TRX (TRX mut, C32S/C35S) a comparable degree (Figures 5G–H). We hypothesized that reducing C41 via TRX WT, but not via TRX mut, would further increase the fraction of p21^high^ cells in G2 phase. Indeed, following Nutlin-3 treatment (1 µM, 24 h) TRX WT, but not TRX mut cells increased p21-GFP levels in G2-phase, mimicking the C41S phenotype (Figure 5I, S5G). No effect was observed in C41S cells (Figure 5J, S5H), indicating that reducing C41 alone is sufficient to stabilize p21 in G2. Of note, expressing VHH increased GFP fluorescence in all cells, likely due to the known GFP-enhancing effects of this nanobody^53^.

To assess functional consequences, we synchronized cells in G1 via palbociclib and released them into S phase with concurrent TRX expression and Nutlin-3 treatment. Live-cell imaging showed that 76% of TRX mut-expressing cells progressed to mitosis within 24 h, compared to only 67% of TRX WT-expressing cells (Figure S5I), suggesting increased p21-dependent G2 arrest^54^.

To test the reciprocal, thus whether increasing C41 oxidation reduces p21 levels, we fused VHH4 to D-amino acid oxidase (DAAO), which generates H₂O₂ upon D-alanine addition^55^ (Figure 5K). Tetracyline-induced expression of VHH4–DAAO in WT and C41S cells to an equal degree (Figure 5L), followed by Nutlin-3 (5 µM, 3 h) and D-alanine (40 mM, 3 h) treatment significantly reduced p21^high^ G2 cells in WT but not C41S cells (Figures 5M–N, S5J–K). This indicates that oxidizing C41 alone is sufficient to reduce p21 levels in G2.

Taken together, C41 redox state controls p21 stability. Reducing C41 prolongs p21 half-life and increases responsiveness to stress-induced expression. Modulating p21’s redox state by localized expression of TRX or DAAO alters p21 levels only in WT cells, confirming that C41 oxidation is sufficient to govern p21 turnover in G2.

### Non-Oxidizable C41 Primes Cells for Cell Cycle Exit into Senescence

Given the increased sensitivity of p21-GFP C41S cells to stress-induced p21 accumulation, and the role of p21 in regulating proliferation versus cell cycle exit^19–21^, we assessed their long-term response to ionizing radiation. Cells were exposed to increasing doses of radiation, and responses were evaluated after six days. C41S cells showed increased dose-dependent p21 accumulation (Figure 6A) and reduced pRB S807/811 staining (Figure 6B) compared to WT, consistent with enhanced cell cycle exit (Figure 6B)^44^. To determine whether these cells entered senescence we measured their nuclear size relative to control (0Gy) and observed that C41S cells show increased nuclear size compared to WT (Figure 6C)^56^. Subsequently, we also measured β-galactosidase activity as a well-established senescence readout (Figures 6C)^57^. While basal β-galactosidase activity was comparable between WT and C41S cells, 6 Gy irradiation markedly increased staining in C41S cells (Figure 6D). Crystal violet staining after 10 days confirmed reduced proliferation in irradiated C41S cells, indicating irreversible growth arrest and senescence (Figures 6E–F).

**Figure 6.**
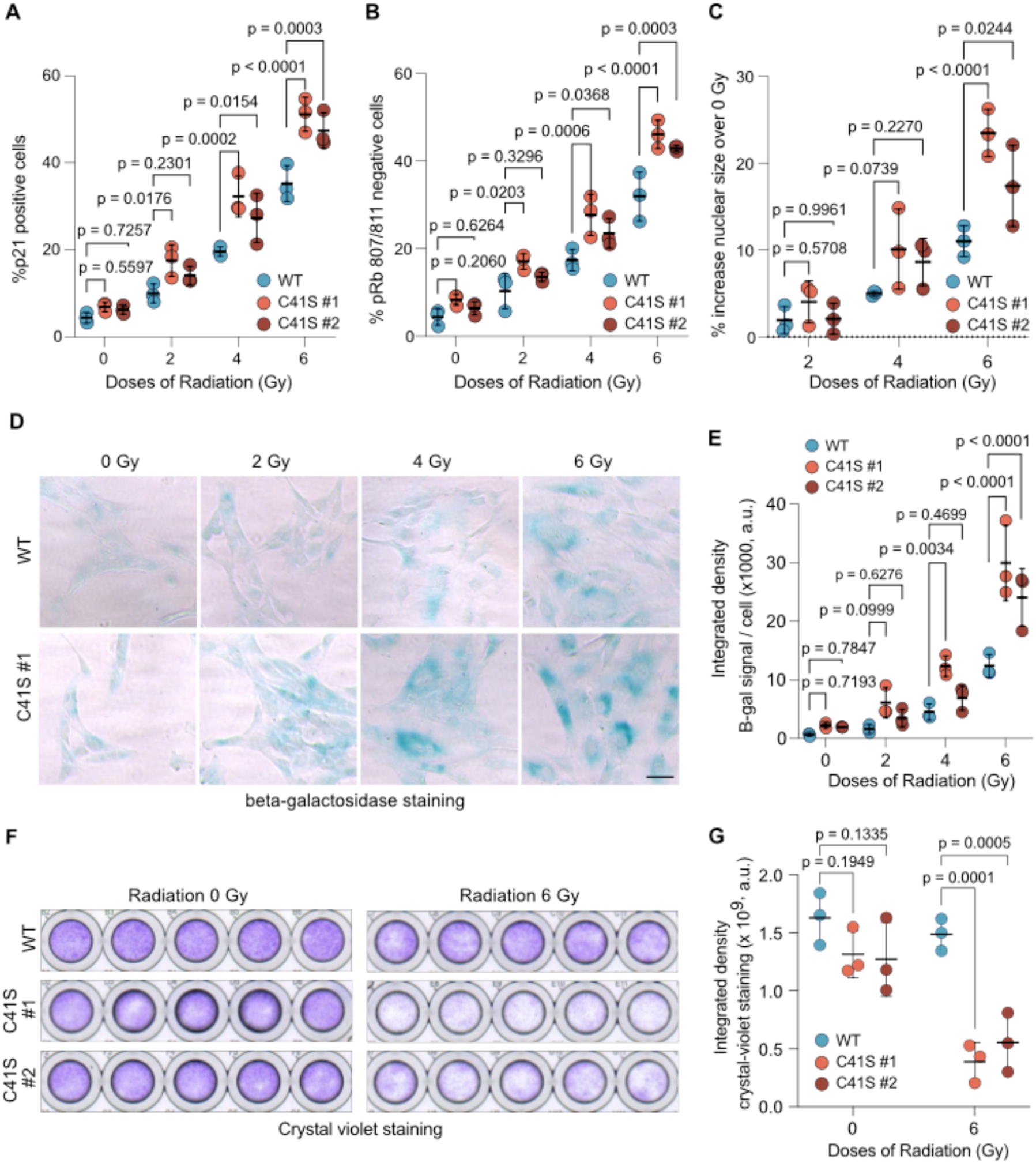
Non-Oxidizable C41 Primes Cells for Cell Cycle Exit into Senescence. (**A-C**) Quantification of the percentage of p21-GFP–positive cells (A), pRB S807/811– negative cells (B), and the percentage increase in nuclear size relative to 0 Gy (C) in RPE-1 p21-GFP WT and C41S cells, measured by fluorescence imaging (A) or immunostaining (B, C) six days after irradiation with the indicated doses. Statistical significance by two-way ANOVA (mean ± SD; n = 3). (**D**) Representative images of β-galactosidase staining in RPE-1 p21-GFP WT and C41S cells six days after irradiation at the indicated doses. Scale bar, 50 µm. (**E**) Scatter plot showing the integrated β-galactosidase staining density per cell from (D). Statistical significance by to two-way ANOVA (mean ± SD; n = 3). (**F**) Crystal violet staining of RPE-1 p21-GFP WT and C41S cells ten days after irradiation with 0 or 6 Gy. (**G**) Quantification of crystal violet staining from (F). Statistical significance by two-way ANOVA (mean ± SD; n = 3).

We conclude that preventing C41 oxidation sensitizes cells to stress-induced p21 accumulation and promotes irreversible cell cycle exit into senescence, highlighting C41’s redox regulation as a key determinant of cell fate under genotoxic stress.

## DISCUSSION

Fluctuating ROS levels in proliferating cells, driven by NADPH oxidases and mitochondrial metabolism, have long been thought to affect thiol oxidation. Early studies noted mitosis-specific thiol staining^8,58^, and recent work showed sulfenic acid accumulation from G1 to G2/M^7,27^ (Figure S1). However, specific redox targets and their functions remained poorly defined.

### Cell Cycle-Resolved Dynamics of S-Sulfenylation

We identified 1,725 sulfenylated cysteines on 1,166 proteins across the cell cycle with 2-hour resolution. Despite flow cytometry showing increasing sulfenic acids from G1-S-G2/M^7,27^, proteome-normalized sulfenylation remained stable (Figure 1). Hence, changes observed by flow cytometry largely reflect changes in protein abundance but not cysteine oxidation. Still, ∼13% of sites exhibited cell cycle-specific dynamics and were enriched in cytoskeletal and cell cycle proteins including IQGAP1, CFL1 and PCM1 (Figure 1, Table S1). This suggests that S-sulfenylation plays a role in cortical changes, mitotic spindle dynamics^59,60^ and centrosome function^61^.

Such dynamics likely result from spatial proximity to ROS sources (e.g., mitochondria, NADPH oxidases^62,63^), and depend on local cysteine environment, peroxiredoxin relays^64^, or redox–phosphorylation crosstalk^65^. Subcellular oxidation domains^66^ may explain heterogeneous sulfenylation, even within the same protein (Table S1).

### Oxidation at C41 Controls p21 Stability

Among cell cycle proteins, p21 was oxidized at C41 in G2 phase with stoichiometries of 2–34% (Figure 2), likely underestimated due to rapid degradation and lack of proteasome inhibition during cell lysis. Modeling based on measured half-lives indicates that the observed average oxidation (12%) is sufficient to rapidly turn over the total p21 pool during G2, assuming a ∼30 min half-life for fully oxidized C41 (Figure S6), consistent with short-lived cell cycle proteins, including p21^48^. Structural analysis of the p21-CDK4-cyclin D trimer placed C41 near CDK4’s C78, a cysteine conserved also in CDK6. Despite CDK4 C78 being oxidized (Table S1), we found no evidence for intermolecular disulfide bond formation between p21 and CDK4. *In vitro*, p21 peptides form intramolecular disulfides (C13–C18, C34–C41) upon H₂O₂ exposure, and overexpression of p21-C18S induces arrest^67^. However, we detected no C18 oxidation in cells, only minor C13 oxidation (4%) in G2 (Figure 2), and BTD proteomics confirmed C41 as the main oxidation site in RPE-1 cells (Figure 1, Table S1). Two findings support that C41 oxidation is sufficient to regulate p21 stability in G2: (i) CRISPR-Cas9-mediated C41 mutagenesis prolonged p21 half-life, inhibited CDK2, and increased steady-state p21 levels in RPE-1 and MCF10A cells (Figures 3, 4); (ii) targeted manipulation of p21 redox state via TRX or DAAO altered p21 levels in WT but not C41S cells, increasing or decreasing p21 in G2, respectively (Figure 5).

### Redox-Regulated Feedback Loop between p21 and CDK2

Our data suggest that reducing C41 enhances p21 binding to CDK2, while oxidizing favors CDK4 interaction, although causality between these effects remains unresolved (Figure 4). IUPred3 analysis predicts that C41 oxidation could alter p21 conformation (Figure 2), potentially weakening CDK2 binding. Increased CDK2 binding by reduced p21 enhances CDK2 inhibition, thus impairing CDK2-mediated S130 phosphorylation, a prerequisite for proteasomal degradation of p21 by SCF-SKP2 in G2 (Figure 4). Accordingly, C41S stabilizes p21 and inhibits CDK2, while the phospho-mimetic S130D mutation restores degradation (Figure 4K). Since p21 phosphorylation relies primarily on the cyclin A interaction, independent of the CDK motif^24,25^, we cannot exclude a role for CDK1-cyclin A in p21 degradation, although cyclin A predominantly associates with CDK2 in G2^68^.

Thus, C41 functions as a redox-sensitive switch tuning the p21-CDK2 feedback loop to restrict p21 inheritance and prevent premature cell cycle exit and senescence (Figure 7). Increased p21 oxidation in G2 may be driven by cell cycle-dependent mitochondrial ROS, as shown for CDK2^9^, or by local ROS generated at DNA damage sites^69^ to support continued proliferation following repair.

**Figure 7.**
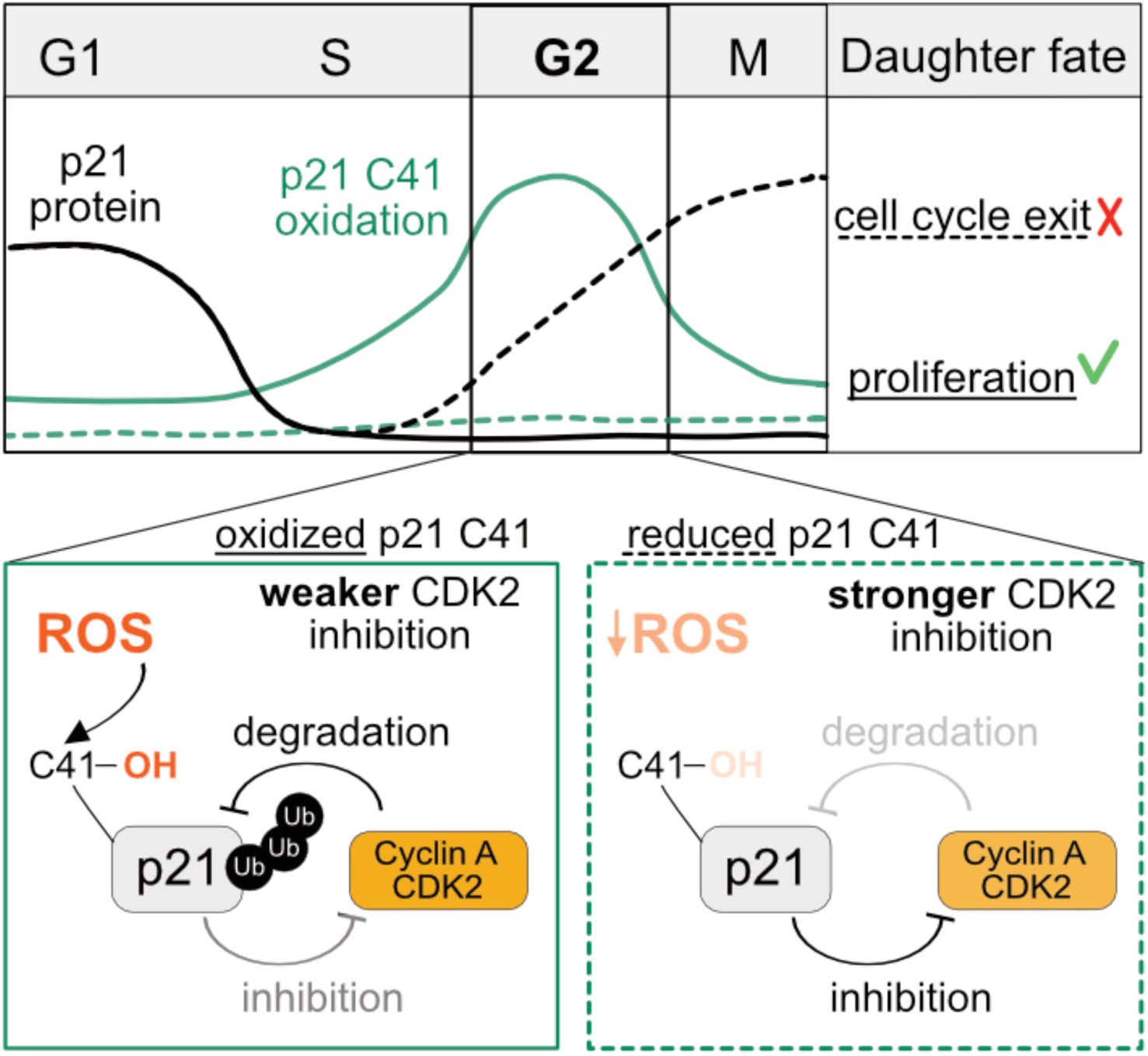
Model of a Redox-Regulated Double-Negative Feedback Between p21 and CDK2. Model illustrating how C41 oxidation regulates the proliferation–cell cycle exit decision by modulating the feedback between p21 and CDK2. Top: Oscillation of total p21 protein levels (black) and C41 oxidation (green) across the cell cycle. Reduced C41 oxidation (dashed green) during G2 slows p21 degradation, resulting in elevated p21 levels (dashed black) inherited by daughter cells, promoting cell cycle exit. Bottom: C41-oxidized p21 is a weaker inhibitor of CDK2–cyclin A, allowing enhanced CDK2-mediated phosphorylation of p21. This phosphorylation creates a phospho-degron recognized by the SCF–SKP2 ubiquitin ligase, leading to proteasomal degradation. In contrast, p21 with reduced C41 more effectively inhibits CDK2–cyclin A, reducing phosphorylation and stabilizing the protein.

### Concluding Remarks

Under normal conditions, only a subset of cells induces p21 in G2 phase^21^, explaining the mild proliferation defects of C41 oxidation-deficient mutants (Figure 3). However, under genotoxic stress or in competitive tumor environments, failure to oxidize C41 becomes a selective disadvantage, halting cell division (Figures 5-6). Targeting C41 oxidation may thus offer a therapeutic strategy for hypoxic tumors, where dormant, p21^high^ cancer cells resist treatment^70^. Tumor re-oxygenation^71^ or inhibition of antioxidant enzymes could enhance C41 oxidation, promote p21 degradation^72^, and sensitize cells to radio- and chemotherapy.

### Limitations of the study

BTD selectively labels sulfenic acids in living cells, avoiding false positives that can arise from atmospheric exposure during sample preparation with differential alkylation– reduction methods. However, BTD has limitations: rapidly converting sulfenic acids (e.g., to disulfides or further oxidized species) may be underrepresented, while long-lived sulfenic acids could be overrepresented. BTD’s covalent binding may also affect protein function; to minimize this, labeling was restricted to one hour as established previously^12^. To identify canonical cell cycle-dependent sulfenylation sites, we correlated individual oxidation events with model profiles showing distinct peaks or declines in G1, S, or G2/M. This approach excluded complex or multiphasic oxidation dynamics (Table S1), whose biological relevance, as illustrated by p21, requires detailed individual analysis.

Our functional experiments focused on modulating the redox state of p21 in living cells using the C41S mutation or localized expression of antioxidant and ROS-generating enzymes (Figure 5). However, there is currently no method to stably and selectively mimic sulfenylation in cells. Future studies will be needed to determine whether sulfenic acid formation at C41 alone is sufficient to alter p21-CDK interactions, or whether secondary modifications such as disulfide formation or metabolite conjugation (e.g., to glutathione or acetyl-CoA) also contribute.

## Resource Availability Lead Contact

Requests for further information and resources should be directed to and will be fulfilled by the lead contact, Jörg Mansfeld.

## Materials Availability

Cell lines and custom analysis scripts are available from the lead contact upon request. Plasmids have been submitted to Addgene.

## Data and Code Availability

Mass spectrometry data have been deposited with the ProteomeXchange Consortium 73§1 (http://www.ebi.ac.uk/pride) under accession numbers PXD054383 and PXD062970 and are publicly available as of the date of publication. Original images have been deposited to Mendeley Data and are available at Mendeley Data: https://doi.org/10.17632/rffhch544x.1

-This paper does not report original code.

-Any additional information required to reanalyze the data reported in this study is available from the lead contact upon request.

## Acknowledgments

We thank Kate Carroll for providing BTD; Alexis Barr for experimental advice, p21-GFP and p21-/- cells; Basil Greber and Lucy Dan for recombinant CDK2–cyclin A; Alex Radzisheuskaya for advice on NGS; Luca Cirillo for support during revisions; and Stefano Pietra for critical reading of the manuscript. We are also grateful to Christian Zierhut, Max Douglas, and Alex Radzisheuskaya for feedback on the manuscript, and to their labs, as well as past and present members of the Mansfeld lab, for technical assistance and valuable discussions. We acknowledge the support of Kai Betteridge and the Light Microscopy and Flow Cytometry Facilities at Chelsea (ICR) and BIOTEC (TU Dresden). J.V. was a member of the Dresden Biomedicine and Bioengineering PhD program. J.K.L. receives funding from the ICR/RM CRUK RadNet Centre of Excellence (grant: A28724). This work was supported by a Cancer Research UK Senior Cancer Research Fellowship to J.M. (RCCSCF-Nov22/100001) and by the European Research Council (ERC) under the EU Horizon 2020 program (REDOXCYCLE, grant 680042 to J.M.; ASYMMEM, grant 758334 to A.N.). T.I.R. and J.S.C. are funded by the CRUK Centre grant C309/A25144, and A.N. receives additional support from the VolkswagenStiftung within the Life? initiative.

## Author Contributions

Conceptualization, J.V. and J.M.; Methodology, J.V., T.I.R., L.Y., J.K.L., D.C., J.C. and J.M; Investigation, J.V., T.I.R., L.Y., J.K.L., D.C., and J.M; Resources, K.B. and A.N.; Mathematical modelling, T.Z. and I.G.; Writing, J.V. and J.M.; Funding acquisition I.G., A.N., J.C., and J.M.; Supervision, I.G., A.N., J.C. and J.M.

## Declaration of Interests

The authors declare no competing interests.

## Declaration of Generative AI and AI-assisted Technologies in the Writing

During the preparation of this work the J.M. and J.V. used ChatGPT 4.o (OpenAI) to correct written text for readability and language. Afterwards J.M. and J.V. reviewed and edited the content as needed and take full responsibility for the content of the publication.

## STAR Methods

### Experimental Model and Study Participant Details

#### Cell Culture and Cell Lines

All cell lines were maintained under standard mammalian cell culture conditions at 37 °C and 5% CO_2_, either at atmospheric (21%) or physiological (6.3%) O_2_ levels, as indicated in the figures. Cells were routinely tested for mycoplasma contamination. The parental hTERT RPE-1 FRT/TR cell line (RRID:CVCL_VP32), used for mass spectrometry, was kindly provided by Jonathon Pines (ICR, London). hTERT RPE-1 FRT/TR mRuby-PCNA + p21-GFP and RPE-1 FRT/TR mRuby-PCNA + p21⁻/⁻ lines used in p21 experiments were gifts from Alexis Barr (Imperial College London). RPE-1 cells were cultured in DMEM/F12 supplemented with 10% FBS, 1% penicillin-streptomycin, 1% GlutaMAX, 0.5 µg/mL amphotericin B, and 0.26% sodium bicarbonate. For experiments at 6.3% O_2_, cells were adapted for ≥5 days in an InvivO2 500 workstation (Baker Ruskinn) and used before day 12 of culture. All media and reagents were equilibrated for ≥2 hours at 6.3% O₂ prior to use. The parental MCF10A line with endogenously tagged mRuby-PCNA, used to generate p21-C41S knock-in clones, was previously established in our laboratory^45^. MCF10A cells were maintained in DMEM/F12 (Invitrogen) supplemented with 5% horse serum, 1% penicillin-streptomycin, 20 ng/mL EGF, 0.5 mg/mL hydrocortisone, 10 µg/mL insulin, and 100 ng/mL cholera toxin. Tetracycline-inducible HA-VHH4-TRX RPE-1 FRT/TR cell lines were generated by electroporating 1–2 × 10^6^ cells in 100 µL tips with plasmids PL4 and PL5 (plus PL6 at a 1:3 ratio) using the Neon Transfection System (2 pulses, 1350 V, 20 ms). After 4 days, cells were expanded and selected with 500 µg/mL neomycin for 14–21 days. Positive clones were identified by anti-HA immunostaining after induction with 100 µg/mL tetracycline.

### Method Details

#### Expression and Purification of Recombinant p21

Full-length p21-WT, p21-C41S, and p21-C41A, each C-terminally tagged with a Twin-Strep tag (IBA), were expressed in *E. coli* BL21 Rosetta (DE3) cells. Protein expression was induced with 0.5 mM IPTG for 4 h at 21 °C. Bacterial pellets were harvested and stored at –80 °C until purification. For purification, frozen pellets were resuspended in lysis buffer (450 mM NaCl, 10% glycerol, 1 mM EDTA, 5 mM β-mercaptoethanol, 1 mM PMSF), lysed by sonication on ice (Sonics Vibra Cell; 40% amplitude, 10 min total; 5 s on/off pulses), and cleared by centrifugation (1 h at 40,000 g, 4 °C). The supernatant was incubated in batch with Strep-Tactin XT resin (IBA) pre-equilibrated in lysis buffer (1 mL resin per prep), followed by 50 column volumes of lysis buffer for washing. Bound p21-StrepII was eluted using BXT buffer (IBA) supplemented with 10% glycerol, 1 mM TCEP, and 450 mM NaCl. Eluates were concentrated using VIVASPIN 6 spin columns (MWCO 5,000; <1 mg/mL), flash frozen in liquid nitrogen, and stored at –80 °C.

#### In Vitro Kinase Assay

Recombinant p21-WT-StrepII, C41S-StrepII, or C41A-StrepII proteins were incubated with recombinant CDK2–cyclin A2 (25 ng per reaction) in kinase buffer (30 mM HEPES, 100 mM NaCl, 10 mM MgCl₂, 5 mM DTT) at 4 °C for 30 min with continuous shaking. Subsequently, recombinant RB protein (aa 773–928) (100 ng per reaction) and ATP (final concentration: 1 mM) were added. For the -ATP control, RB was added but ATP was omitted. Reactions were incubated for 15 min at 30 °C with shaking and terminated by addition of 4x NuPAGE sample buffer followed by boiling at 95 °C for 5 min. DTT was then added to a final concentration of 100 mM prior to SDS-PAGE and Western blotting. Related to Figure S3.

### Crispr-Cas9-Engineering and NGS Sequencing

CRISPR-Cas9-mediated gene editing was performed using ouabain selection as described previously^40^ based on a plasmid kindly provided by Yannick Doyon. An additional U6-gRNA cassette with BsmBI sites was inserted to allow cloning of two gRNAs targeting CDKN1A/p21 (plasmid PL3). Guide RNA against ATP1A1 was used as previously established^40^ (RD1, Table S2) and guide RNAs against CDKN1A were selected using the CRISPick tool (Broad Institute; gRNAs RD2, RD3, Table S2). Phosphorothioate-modified single-stranded DNA oligonucleotides (ssODNs) serving as HDR templates for ATP1A1^40^ (RD4) and p21 (RD5) were synthesized by IDT (Table S2). To facilitate screening of C41S integrants, a silent XhoI site was included in RD5 at wobble positions (Figure S3A).

For generation of p21-GFP C41S clones in RPE-1 cells, 300,000 p21-GFP cells were electroporated using the Neon Transfection System (10 µL tip; 2 pulses, 1350 V, 20 ms) with 500 ng PL3 plasmid, 2 pmol ATP1A1 ssODN (RD4), and 6 pmol p21 ssODN (RD5). MCF10A cells (200,000) were nucleofected using the Amaxa 4D Nucleofector X Unit (program DS-138; SE Cell Line Kit) with the same nucleic acid mix as above. After 3–4 days, cells were plated onto 15 cm dishes and treated with 0.25 µM ouabain. Single-cell clones were isolated 7–12 days later and expanded. Genomic DNA was extracted using the EZNA Tissue DNA kit (Omega Bio-Tek), and edited loci were amplified using primers listed in Table S2. Amplicons for restriction digestion (PCR1 + PCR2) and next-generation sequencing (PCR3-8 + PCR9) were generated. For amplicon-NGS (GeneWiz, Azenta), six-nucleotide barcodes were added to allow multiplexing of WT and C41S clones. Data were analyzed using CRISPResso2^73^ with the following parameters: CRISPResso --fastq_r1 1_R1_001.fastq.gz --fastq_r2 1_R2_001.fastq.gz --amplicon_seq seq1, seq2,seq3,seq4,seq5,seq6 --amplicon_name cl1,cl2,cl3,cl4,cl5,cl6, --guide_seq GCGACTGTGATGCGCTAAT --trim_sequences --trimmomatic_options_string ILLUMINACLIP:TruSeq3-PE.fa:2:30:10:2:True --min_average_read_quality 30 -- max_paired_end_reads_overlap 150 -qwc 112-132 -n sample_1; Allele frequency tables were used to determine genotypes (Figure S3) and to calculate clone abundance in cell competition assays (Figure 3). Related to Figures 3 and S3.

### BTD

BTD was used to label sulfenic acids in living cells, as previously described^12,26^. BTD for initial testing was kindly provided by Kate Carroll (UF Scripps Biomedical Research, USA). BTD used for mass spectrometry, flow cytometry, subcellular localization, and pull-down experiments was synthesized according to the protocol in^26^ by André Nadler and Kirstin Bohlig (MPI-CBG, Dresden, Germany). Lyophilized BTD was dissolved in anhydrous 100% DMSO at a concentration of 200 mM, aliquoted, and dried under vacuum. For each experiment, individual aliquots were freshly reconstituted in 100% DMSO and vortexed thoroughly until fully dissolved. To maintain probe integrity, aliquots were frozen only once after resuspension in DMSO. Related to Figures 1, 2, and S1.

### Cell Cycle Synchronization and Cell Treatments

For synchronization, RPE-1 cells were seeded at low density (<8,000 cells/cm²) and treated with 150 nM palbociclib for 24 hours. Cells were then washed 2–3 times with warm PBS and released into fresh culture medium. Samples were collected at the indicated time points post-release. For p21 oxidation stoichiometry and half-life measurements, cells were synchronized in G2 phase by releasing them from palbociclib for 11 hours. Cycloheximide (CHX, 50 µg/mL) was added after release for the indicated durations. For half-life experiments with tetracycline-inducible p21-HA cells asynchronous cultures were treated with 50 µg/mL CHX for the indicated time periods. To induce p53, cells were either treated with Nutlin-3 at the indicated concentrations and durations or irradiated using a Gulmay X-ray generator with MP1 controller at the specified doses (Gy). For live-cell imaging of TRX-expressing cells, synchronization was performed as above. Seven and a half hours post-release, Nutlin-3 (1 µM final concentration) and tetracycline (see Figure S5I) were added to the medium. Related to Figures 1, 2, 4, 5, S1, S4, and S5.

### Doubling-time Analysis

To assess the doubling time of cells, a defined number of cells were seeded into 6-well dishes and the time until the next splitting precisely noted. Before cells reached confluency (usually 3-4 days) they were trypsinised, collected in media, fixed in 4% paraformaldehyde (PFA) for 10 minutes, pelleted, and resuspended in PBS for cell counting using an automated cell counter (Countess, Thermo Fisher). Then the doubling was calculated according to: *Doubling Time = [T × (ln2)] / [ln (Xe / Xb)].* With T = time, Xe = number of cells at the end, Xb= number of cells at the beginning. Related to Figure 3.

### Growth-Competition and Reversion

p21-WT and p21-C41S (#1 or #2) cells were mixed roughly 50:50, seeded into dishes, and a sample was taken for later amplicon-NGS analysis (day 0). The mixed cell population was passaged regularly before cells reached confluency and at day 12 and day 20 samples were taken for NGS analysis. For amplicon-NGS genomic DNA was extracted and a short genomic region around C41 of CDK1NA (p21) was amplified using primers indicated in Table S2. The resulting PCR product was gel-purified and sent for amplicon-NGS by GeneWiz (Azenta Life Science). Different timepoint were analysed in parallel in Crispresso2^73^ using bar coded primers as detailed above. The alleles frequency table was used to collect the precise number of reads corresponding to the WT or C41S sequence and used to express the ratio of C41S/WT of day 12 and 20 relative to day 0.

To revert p21-GFP C41S or p21-GFP WT, Cas9 was electroporated as a ribonucleoprotein (RNP) complex with either RD3 (targeting WT) or RD6 (targeting C41S) (see Table S2), along with a single-stranded oligodeoxynucleotide (ssODN) repair template encoding the WT C41 sequence (RD7). To distinguish reverted from uncut endogenous WT sequences, the repair template contained silent wobble mutations. TracerRNAs and gRNAs were mixed in a 1:1 ratio and mRNPs assembled with a 1:1.2 ratio of NLS-Cas9: gRNA complex. 300,000 RPE-1 p21-WT-GFP or p21-C41S-GFP cells were electroporated using the Neon Transfection System (10 µl tip, two pulses at 1350 V for 20 ms) with 30 pmol mRNPs and 0.2 pmol ssODN repair template. Cells were collected at day 1 and day 4 post-electroporation for genomic DNA extraction and PCR amplification of the C41 locus (using barcoded primers in Table S2). Amplicons were purified using Mag-Bind DNA beads, submitted for amplicon-NGS (GeneWiz), and analyzed with CRISPresso2^73^ as described. Allele frequency tables were used to determine the proportion of parental (WT or C41S) and successfully reverted sequences. Related to Figure 3.

### β-Galactosidase Staining

Senescence-associated β-galactosidase activity was assessed using a Senescence β-Galactosidase Staining Kit (Cell Signaling Technology) according to the manufacturer’s instructions. Briefly, p21-GFP WT and C41S (#1 and #2) cells were irradiated with the indicated doses and passaged once 2–3 days later to avoid confluency. Six days post-irradiation, viable cells were counted using Trypan Blue exclusion and seeded into clear 96-well plates. The following day, cells were washed twice with PBS and fixed for 15 min using the provided fixative solution. After two additional PBS washes, cells were incubated overnight at 37°C (dry incubator, no CO₂) with β-galactosidase staining solution (pH 5.9–6.1). Cells were then washed twice with PBS and incubated with 1 µg/mL DAPI for 10 min to stain nuclei, followed by two final PBS washes before imaging. Related to Figure 6.

### BTD Staining for Fluorescence Microscopy

RPE-1 cells were seeded in IBIDI microscopy slides and incubated with 1 mM BTD or 0.5% DMSO (vehicle control) for 1 hour in serum-free medium. Cells were washed twice with PBS and fixed in 4% paraformaldehyde (PFA)/PBS containing 5 mM iodoacetamide (IAA) to block reduced thiols. After two PBS washes, cells were permeabilized with −20°C methanol (90%) for 15 min, washed again in PBS, and blocked in 5% BSA/PBS for 30 min at room temperature on a horizontal shaker. BTD-labelled proteins were conjugated to azide-488 via CuAAC (click chemistry). The click reaction mix contained 1 mM CuSO_4_, 0.1 mM Tris[(1-benzyl-1H-1,2,3-triazol-4-yl)methyl]amin) (TBTA), 1 mM sodium ascorbate, and 0.1 nmol azide-488. Cells were incubated with the mix for 1 hour in the dark. After removal of the click solution, cells were washed three times in PBS containing 20 mM EDTA and 0.02% Tween, stained with 1 µg/mL DAPI for 10 min, washed once more, and imaged. Related to Figure S1.

### Crystal-Violet Staining

p21-GFP WT and C41S (#1 and #2) cells were irradiated with the indicated doses (Gy) and passaged once 2–3 days later to prevent confluency. Six days after irradiation, viable cells were counted by Trypan Blue exclusion, and exactly 500 living cells were seeded per well for each condition. After four additional days of culture (10 days total), cells were harvested once p21-GFP WT cells reached confluency. No floating cells were observed during the experiment. Cells were washed twice with PBS and fixed in ice-cold 90% methanol for 10 min. After removing the fixative, cells were stained with 0.1% crystal violet in 25% methanol for 10 min, followed by four washes with distilled water. Plates were dried completely and stored protected from light prior to imaging. Related to Figure 6.

### Immunofluorescence Staining

To detect pRb (S807/811) and γH2AX, cells were seeded into black 96-well plates with a clear bottom, cultured for 2–3 days, washed twice in PBS, and fixed with 4% PFA/PBS for 15 min at RT on a horizontal shaker. For normoxic experiments, fixation was performed in an InvivO2 500 workstation (6.3% O₂) before processing at atmospheric oxygen. For p21-HA degradation experiments (Figure 4), cells were fixed directly in 4% PFA in culture medium inside the incubator. After 15 min, PFA was replaced with PBS until all timepoints were collected. Cells were then washed twice in PBS, permeabilized with 0.2% Triton X-100 + 0.01% SDS in PBS for 10 min, and blocked in 2% BSA in PBS/0.1% Triton X-100 for 30–60 min. Primary antibodies were added in blocking buffer and incubated for 1 hour at RT or overnight at 4°C. After two PBS washes, secondary antibodies were applied for 1 hour at RT, followed by three PBS washes, DNA staining with 1 µg/mL Hoechst-33342 for 15 min, and two final PBS washes before imaging. Related to Figures 3, 5, 6, S1, S3, and S5.

### Flow Cytometry

For flow cytometry cells were trypsinised, washed once in PBS, fixed in ice-cold 70% ethanol, washed in PBS and resuspended in 50µg/ml propidium iodide (PI)-staining solution supplemented with 100µg/ml RNAse. Samples were analysed on an LSRII– analyser (BD) and 10.000 events per sample collected to create cell cycle profiles indicated in Figure 1B. For cell cycle sulfenic acids analysis, cells were labelled with BTD for 30 minutes in cell culture media containing 1% serum, then washed twice, trypsinised, and fixed with 4% PFA/PBS supplemented with 5mM IAA. Then cells were permeabilized with 90% ice-cold methanol and blocked for 30 minutes in 5% BSA-PBS. Click-reactions with 3 million cells containing 2nmol azide-488 were performed in click buffer (1mM CuSO4, 0.1mM TBTA, 1mM sodium ascorbate) for 1 hour on a rotation-wheel in the dark. Afterwards, cells were washed 3x in PBS containing 20mM EDTA, stained with 1µg/ml DAPI, washed 3x in PBS and 50.000 events were analysed on an LSRII–analyser (BD). Data analyses were performed with FlowJo TM (Becton Dickinson & Company (BD) 2023, version: 10.9) Related to Figures 1 and S1.

### SDS-Page and Western blot

Total cell lysates were prepared by washing cells twice in PBS and lysing directly in 1x NuPAGE LDS Sample Buffer supplemented with 100 mM DTT, unless otherwise indicated. Samples were sonicated using a tube sonicator and boiled for 5 min at 95 °C. Proteins were separated on 4–12% NuPAGE Bis-Tris gels using MES or MOPS buffer and transferred to Immobilon-FL PVDF membranes via wet transfer (20% ethanol/MOPS) using Bio-Rad Mini Trans-Blot Cells or a Criterion Blotter (400 mA, ≥1.5 h). Membranes were blocked for 1 hour at RT in 5% dry milk in PBS/0.02% Tween-20. For phospho-specific antibodies, blocking and primary antibody incubation were performed in 4% soy milk. Primary antibodies were incubated overnight at 4 °C, followed by 3× washing in PBS/0.02% Tween-20 and 1 hour incubation with secondary antibodies at RT. Detection was performed using either fluorescent secondary antibodies on an Odyssey system (Li-COR) or HRP-conjugated antibodies with Luminata Forte or SuperSignal West Femto substrate and imaged on an Azure Biosystems chemiluminescence system. Related to Figures 2, 4, 5, S3, and S4.

### Detection of S-sulfenylated p21 by Western blot

Asynchronously growing RPE-1 cells were labelled for 2 h with 1 mM BTD or 0.5% DMSO in serum-free media, washed twice in PBS containing 100 U/mL catalase, trypsinized, pelleted, and lysed in buffer M (50 mM HEPES pH 8, 150 mM NaCl, 1% SDS) supplemented with cOmplete™ protease/phosphatase inhibitors, 200 U/mL catalase, and 10 mM TCEP. Lysates were sonicated and incubated for 30 min at 25 °C in the dark on a horizontal shaker. Following reduction with TCEP, 40 mM IAA was added and incubated for 30 min under the same conditions. Lysates were cleared by centrifugation (14,000 g, 15 min, 20 °C), and protein concentration determined via BSA assay. At least 1 mg protein was used for CuAAC click chemistry with azide-biotin (1 mM CuSO_4_, 0.1 mM TBTA, 1 mM sodium ascorbate, 10 nmol azide-biotin) for 1 hour at 25 °C in the dark. Reactions were quenched with 20 mM EDTA, and proteins precipitated via chloroform/methanol (4:1:3 MeOH:CHCl_3_:H_2_O). Pellets were resuspended in buffer S (50 mM HEPES pH 8, 0.5% SDS, 0.5 M urea), incubated until dissolved (∼15 min), diluted 1:1 with H_2_O, and incubated with streptavidin sepharose overnight at 4 °C. Beads were washed 4x with wash buffer (50 mM HEPES pH 8, 4 M urea, 0.5% SDS), transferred to fresh tubes, washed twice more with wash buffer and once with PBS. Bound proteins were eluted by boiling in LDS sample buffer containing 2.5 mM biotin (15 min, 95 °C), resolved by SDS-PAGE, and analyzed by Western blot. Related to Figure 2.

### p21-GFP Pull Down

Recombinant GFP nanobody VHH4^38,39^ was covalently coupled to carboxylic acid beads (Invitrogen) according to manufacturer instructions (GFP-binder beads). To assess p21-GFP interactions with CDK-cyclin complexes, asynchronous or G2-synchronized p21-GFP WT and p21-GFP C41S (#1) RPE-1 cells were used. Where indicated, synchronized cells were treated with 10 µM H_2_O_2_ for 30 min prior to lysis. Cells were lysed on ice for 15 min in buffer P (50 mM Tris pH 8.0, 100 mM NaCl, 2.5 mM MgCl_2_, 5% glycerol, 0.5% Triton X-100) supplemented with cOmplete™ protease/phosphatase inhibitors and, for reducing conditions, 10 mM freshly prepared DTT. Lysates were cleared by centrifugation (14,000 g, 10 min, 4 °C), protein concentrations determined, and ≥500 µg of lysate added per pull-down to GFP-beads pre-equilibrated in buffer P. After 1 hour incubation on a rotator at 4 °C, beads were washed 3× in buffer P and transferred to fresh tubes. Proteins were eluted by boiling in NuPAGE LDS buffer at 65 °C for 10 min. Eluates were supplemented with 100 mM DTT prior to SDS-PAGE and Western blotting. Beads without VHH4 were used as negative controls. Related to Figures 4 and S4.

### Cell Cycle BTD Proteomics

RPE-1 cells were seeded at low density (<8000 cells/cm²) in 15 cm dishes and synchronized into distinct cell cycle phases using palbociclib. Cells were subsequently labelled with 1 mM BTD for 1 hour in serum-free medium (Figure 1A). After labelling, cells were washed twice with PBS supplemented with 100 U/mL catalase to prevent artificial oxidation, trypsinized, collected in PBS/catalase, and flash-frozen in liquid nitrogen. A small aliquot was fixed in ice-cold 70% ethanol for flow cytometry to determine cell cycle distribution (Figure 1B). Frozen pellets were lysed in buffer M (50 mM HEPES pH 8.0, 150 mM NaCl, 1% SDS) supplemented with HALT™ protease/phosphatase inhibitors, 200 U/mL catalase (in 50 mM potassium phosphate, pH 7.0), and 10 mM TCEP. Lysates were sonicated and incubated for 30 min on a horizontal shaker. Proteins were alkylated with 40 mM IAA for 30 min under the same conditions. After centrifugation, 2.5 mg protein per timepoint was subjected to CuAAC-click chemistry using 25 nmol azide-picolyl-Dde-biotin in 1 mM CuSO_4_, 0.1 mM TBTA, and 1 mM sodium ascorbate at 20 °C for 1 hour, protected from light. Reactions were quenched with 20 mM EDTA. Proteins were precipitated using chloroform/methanol (Cl/MeOH), pelleted (4,000 rpm, 15 min), washed with 4 volumes of MeOH, pelleted again, and air-dried at 37 °C. Pellets were resuspended in 100 mM TEAB (pH 8.0), sonicated until dissolved, and digested overnight with trypsin (in 0.5% formic acid) at 37 °C. The following day, insoluble material was removed by centrifugation (13,000 rpm, 5 min, RT), and 15 ng of each digest was reserved for total proteome analysis.

Remaining peptides from each condition were incubated overnight with 100 µL streptavidin sepharose beads per sample. After two washes in TEAB, bound peptides were labelled with TMT-18plex reagents for 1 hour at 25 °C (40 µL of reagent in 100% ACN, final volume 200 µL with 20% ACN). Labelling was quenched with 5% hydroxylamine. Beads were washed twice with TEAB containing 500 mM NaCl and 0.5% SDS, pooled, and subjected to three additional washes in TEAB and two in water. Peptides were eluted by Dde-cleavage using 2% hydrazine at 20 °C for 30 min on a shaker. Eluates were cleared from beads using TopTips, dried, and prepared for MS injection. Peptides used for total proteome analysis were labelled separately following the manufacturer’s TMT-18plex protocol. Related to Figures 1 and S1.

### p21-Oxidation Stoichiometry

RPE-1 cells were synchronized in G2 phase with palbociclib as described above, washed with PBS, trypsinized, quenched in complete medium, washed again in PBS, and split into two equal aliquots for total cysteine and oxidized cysteine measurements. Cells were pelleted and flash-frozen in liquid nitrogen for subsequent analysis. For lysis, cell pellets were resuspended in buffer M2 (100 mM TEAB, pH 8.5, 150 mM NaCl, 1% sodium deoxycholate, 1 mM EDTA, 10 µM neocuproine) supplemented with fresh cOmplete™ protease and phosphatase inhibitors and 200 U/mL catalase. For total cysteine samples, the buffer was immediately supplemented with 10 mM TCEP; for oxidized cysteine samples, 10 mM IAA was added instead. For H_2_O_2_-treated controls (used only as tracer control in MS), cells were lysed in buffer M2 supplemented with 10 or 100 µM H_2_O_2_ and incubated for 30 min at room temperature. Subsequently, catalase (200 U/mL) and either 10 mM TCEP (for total cysteine samples) or 10 mM IAM (for oxidized cysteine samples) were added. All samples were homogenized using a tube sonicator and incubated at 37 °C for 1 hour, protected from light. Protein concentration was determined, and 500 µg of each sample was used for downstream processing. Excess IAA and TCEP were quenched with 10 mM L-cysteine, followed by simultaneous reduction and alkylation with 40 mM iodoacetamidopropionyl (IPM) at 37 °C for 1 hour in the dark. Proteins were incubated with carboxylic acid–functionalized magnetic beads (Invitrogen) crosslinked to the anti-GFP nanobody VHH4 and equilibrated in M2 buffer. After 1 hour of rotation at 4 °C, beads were washed 3 times with buffer M2 supplemented with 500 mM NaCl, transferred to fresh tubes, and washed twice more with buffer M2. Beads were then resuspended in 100 mM TEAB and digested overnight with trypsin (50 ng/µL). The resulting peptides were collected and labelled using TMT-10plex reagents following the manufacturer’s instructions. Related to Figure 2.

### Mass Spectrometry Analysis and Database Search

TMT labelled peptides for total proteome analysis were fractionated with high-pH Reversed-Phase (RP) chromatography using the XBridge C18 column (2.1 × 150 mm, 3.5 μm, Waters) on an UltiMate 3000 HPLC system. Mobile phase A was 0.1% (v/v) ammonium hydroxide and mobile phase B was ACN, 0.1% (v/v) ammonium hydroxide. The peptides were fractionated at 0.2 mL/min with the following gradient: 5 minutes at 5% B, up to 12% B in 3 min, for 32 min gradient to 35% B, gradient to 80% B in 5 min, isocratic for 5 minutes and re-equilibration to 5% B. Fractions were collected every 42 sec SpeedVac dried and eventually pooled into 30 samples for LC-MS analysis. The TMT labelled peptides from p21 pull-downs were fractionated with a 1×100 mm RP column at 70 μL/min and 8 (10plex) or 12 (18plex) fractions were eventually collected for LC-MS analysis.

LC-MS analysis for the BTD and total proteome experiments was performed on an UltiMate 3000 UHPLC system coupled with the Orbitrap Lumos mass spectrometer (Thermo Scientific). Peptides were loaded onto the Acclaim PepMap 100, 100 μm × 2 cm C18, 5 μm, trapping column at flow rate 10 μL/min and analysed with an Acclaim PepMap (75 μm × 50 cm, 2 μm, 100 Å) C18 capillary column connected to a stainless-steel emitter on the EASY-Spray source. Mobile phase A was 0.1% formic acid and mobile phase B was 80% ACN, 0.1% formic acid. Peptides were analysed with a gradient of 5%-38% B in 125 min (BTD runs) or 95 min (total proteome runs). MS scans were acquired in the range of 375-1,500 m/z at a mass resolution of 120,000. Precursors were selected with the top speed mode in 3 sec cycles and isolated for HCD fragmentation at 30k or 50k resolution with 36% collision energy (CE). Targeted precursors were dynamically excluded for 30 or 45 seconds with 7 ppm mass tolerance.

The analysis of the p21 pull-downs (IPM experiment) was performed on an Orbitrap Ascend mass spectrometer (Thermo Scientific) with a Synchronous Precursor Selection (SPS) MS3 method over a 100 min gradient (3%-27% B) on an UltiMate 3000 UHPLC for the TMT10plex runs, or with an SPS Real Time Search (RTS) MS3 method over a 110 min gradient (5%-35% B) for the TMT18plex runs on a Vanquish Neo UHPLC using the nanoE MZ PST BEH130 C18, 1.7 μm, 75 μm × 250 mm analytical column (Waters). MS2 spectra were collected with HCD fragmentation (CE=32%) and ion trap detection whilst the MS3 quantification spectra were collected with HCD fragmentation (CE=55% or 65%) and Orbitrap detection at 45k resolution. Static modifications for RTS were TMTpro16plex at K/n-term (+304.2071), IPM at C (+95.03711) and variable modifications were Deamidated NQ (+0.984) and Oxidation of M (+15.9949) with maximum 1 missed-cleavage and 2 variable modifications per peptide.

The mass spectra were processed in Proteome Discoverer (Thermo Scientific) v2.4 for the total proteome data or v3.0 for the BTD and p21 pull-downs with the SequestHT and Comet (for v3.0 only) search engines for peptide identification and quantification. For the BTD and total proteome experiments the precursor and fragment ion mass tolerances were 20 ppm and 0.02 Da respectively. For the p21 pull-downs MS2 tolerance was 0.5 Da. Spectra were searched for fully tryptic peptides with maximum 2 missed cleavages. TMTpro or TMT6plex at N-terminus/K were selected as static modifications according to each experimental design. Oxidation of M and deamidation of N/Q were selected as dynamic modifications in all searches. Additionally, for the BTD experiments, BTD+DdePicolyAzide_Cleaved (+481.153 Da) and Carbamidomethyl (+57.021 Da) at C were selected as dynamic modifications. Carbamidomethyl at C was set as static for the total proteome data. For the p21 pull-downs, IPM (+95.03711) and Carbamidomethyl at C were added as dynamic modifications. Spectra were searched against reviewed UniProt Homo Sapiens protein entries; peptide confidence was estimated with the Percolator node and peptides were filtered at q-value<0.01 based on target-decoy database search. The reporter ion quantifier node included a TMTpro or TMT6plex quantification method with an integration window tolerance of 15 ppm. Related to Figures 1, 2 and S1.

### Imaging

Automated microscopy was performed using an ImageXpress Micro Confocal system (Molecular Devices) in wide-field mode, equipped with 10×/0.45 NA and 20×/0.7 NA Plan Apo air objectives (Nikon) and a laser-based autofocus module. Excitation and detection were achieved using a Spectra X light engine (Lumencor) and a 16-bit sCMOS camera (Andor). The following filter sets were used: DAPI: Ex 377/54; Dichroic 409; Em 447/60, FITC/GFP: Ex 475/34; Dichroic 506; Em 536/40, TexasRed: Ex 560/32; Dichroic 593; Em 593/40, Cy5: Ex 631/28; Dichroic 660; Em 624/40. All imaging was performed with binning = 1. For live-cell experiments, cells were maintained in a stage-top incubator at 37 °C in a humidified atmosphere with 5% CO₂. Imaging was conducted in phenol red–free DMEM supplemented with 10% FBS, 1% penicillin-streptomycin, 1% Glutamax, 0.5 μg/mL Amphotericin B, and 0.26% sodium bicarbonate. A single position per well was acquired. For immunofluorescence imaging, at least nine positions per well were acquired with binning = 1. All cells were seeded in black 96-well clear-bottom plates (μClear, Greiner Bio-One). For crystal violet–stained cells, imaging was performed using a LI-COR Odyssey imaging system equipped with an RGB-Trans scanning module. For β-galactosidase and DAPI staining, cells were imaged using an EVOS M5000 Imaging System (Thermo Fisher) with an RGB Trans option, which generates color images from interlaced RGB channels. Images were acquired using a 10×/0.30 NA objective. Detection of DAPI and β-galactosidase staining was performed using the tagBFP, GFP, and TexasRed filter cubes. At least two randomly selected positions per well were acquired. For visualization of BTD-labelled sulfenic acids in fixed cells, imaging was performed on a DeltaVision wide-field deconvolution fluorescence microscope (Imsol) equipped with a PCO Edge sCMOS camera. Images were acquired using an Olympus UPlan Apo 20×/0.7 NA objective at binning = 1 or 2. DAPI and Cy5 signals were collected using the following filters: DAPI (381–412 nm), Cy5 (625–646 nm). Related to Figures 3–6, S1, S3–S5.

### Semi-Automated Image Analysis and Immune Fluorescence Staining

IF images acquired with the ImageXpress system were analyzed using automated pipelines developed in the Custom Module Editor of MetaXpress (Molecular Devices). Images were flat-field and background corrected using a top-hat filter. Nuclei were segmented based on Hoechst 33342 staining to generate nuclear masks, which were used to extract fluorescence intensities for p21-GFP, pRb (S807/811), γH2AX, and p21-HA. Positive cells were identified by applying fixed thresholds to flat-field–corrected Cy5 or GFP channels (γH2AX-Cy5, pRb-Cy5, or p21-GFP), based on visual inspection. All conditions and cell lines were processed in parallel using identical thresholds. For single-cell analysis, integrated fluorescence intensities of p21, pRb, and Hoechst 33342 were extracted. Radiation-induced changes in nuclear size were calculated as a percentage relative to untreated controls (0 Gy). CDK2 activity was assessed by computing the cytoplasmic-to-nuclear ratio of GFP signal from the CDK2 activity sensor. A cytoplasmic ring mask was generated by dilating the nuclear mask by five pixels and subtracting the original mask. Cells with cytoplasmic/nuclear GFP ratios outside the range of 0–2.5 were excluded to avoid segmentation artifacts. To assess p21 degradation in p21-HA cells, the 90th percentile of HA intensity in G2-phase cells was measured and background-subtracted using the 90th percentile HA intensity from S-phase cells to correct for antibody background staining. Cell cycle phases (S and G2) were determined based on Hoechst-33342 staining. Related to Figures 3, 4, 5, 6, S3, S4, and S5.

### Image Analysis of β-Galactosidase and Crystal-Violet Staining

Quantification of β-galactosidase staining per cell was performed using a Fiji plugin developed by^57^, enabling automated and unbiased image analysis. This workflow provides a more robust and objective assessment of senescence-associated β-galactosidase (SA-β-Gal) activity compared to manual scoring. Threshold intensity values were defined using cells irradiated with 6 Gy and subsequently applied uniformly to all images. DAPI-stained nuclei were counted simultaneously, and β-galactosidase signal intensities were normalized to the number of nuclei to calculate signal per cell. For crystal-violet staining, images were opened in Fiji (version 2.14.0/1. 54f)^74^ and converted to RGB format. Integrated densities for each entire well were measured, and background signals were subtracted accordingly. Related to Figure 6.

### Semi-automated Image Analysis of Live Cell Imaging

Semi-automated image analysis was implemented using custom ImageJ macros alongside Jython and Python scripts. Background subtraction and illumination correction were carried out using BaSIC^75^. As G0 phase nuclei lack mRuby-PCNA signal, a pseudo-channel combining the p21-GFP and mRuby-PCNA channels was created for segmentation. To address over-segmentation caused by mRuby-PCNA replication foci in S phase, nuclei were segmented using StarDist^76^ on Ilastik^77^ pixel probability maps trained to classify “nuclei” versus “background” in the pseudo-channel. Object tracking across time was performed using the linear assignment problem (LAP) algorithm in TrackMate^78^. Tracks shorter than 24 hours were excluded. Manual curation retained only tracks containing one S/G2 transition and one mitosis. Tracks were terminated if (1) a G1– S transition occurred, (2) the cell exited the field of view, or (3) segmentation/tracking failed. Feature extraction and analysis were conducted using Pandas and Matplotlib in Python. A ±2 frame rolling mean filter was applied to all track features. Curated tracks were in silico–synchronized at the S/G2 transition, defined as the frame with the highest standard deviation of mRuby-PCNA intensity. Mitosis was defined as the frame with the largest object area (typically 2–20 hours after S/G2), reflecting cytoplasmic diffusion of mRuby-PCNA upon nuclear envelope breakdown. G2 timepoint values were calculated as the midpoint between S/G2 and mitosis; G1 measurements were taken 3 hours post-mitosis (Figure S3N). For TRX movies (Figure S5I), only tracks starting in S phase were retained after manual curation. As above, S/G2 transitions were defined by PCNA intensity deviation, and mitosis was marked as the frame with the largest nuclear size change relative to frame n–2. Tracks were excluded if they did not persist at least five times the average G2 duration in unperturbed cells (Figure 3G, 4.78 h) after G2 entry, to prevent misclassification due to tracking errors. Related to Figures 3, S3, and S5.

### Data Visualization

Fluorescence microscopy and scanned images (crystal violet and Western blots) were processed using Fiji (version 2.14.0/1. 54f)^74^ applying identical brightness and contrast settings per channel within each experiment. Cell cycle–resolved p21 sulfenylation was visualized using Cytoscape (version 3.1)^79^. Scatter plots, bar charts, and XY plots were generated with GraphPad Prism (version 10.0). Bivariate density plots and gene ontology (GO) analyses were performed in RStudio (version 2024.04.2+764; R version 4.4.1) using the tidyverse, ggpointdensity, ggplot2, and gprofiler2 packages. For all density plots, the number of cells was equalized between conditions. Single-cell and mean p21 intensities from live-cell imaging were plotted using Python (version 3.11.4).

### Structures and protein conservation

ChimeraX (version 1.6.1)^80^ was used to visualize publicly available structures of p21 bound to CyclinD–CDK4 (PDB: 6p8h^32^) and a model of p21 bound to CyclinA-CDK2^33^. Distances between p21 and Cyclin–CDK complexes were measured using the ChimeraX distance calculation tool. Protein conservation was assessed using ProViz^81^, and multiple sequence alignments of p21 with other CDK inhibitors were generated using the online version of Clustal Omega^82^. Related to Figures 2 and S2.

### Model of p21 oxidation and stability

To estimate the influence of different degradation rates on the overall stability of total p21, the dynamical equilibrium between reduced p21 (*p_red_*) and C41-oxidised p21 (*p_ox_*) was described in terms of ordinary differential equations (ODE) with constant production (rate *b*) and fast interconversion between the species (rates *k_+_* and *k___*). First order decay dynamics were parameterized by different half-lives of reduced p21 (*t_red_*) and C41-oxidised p21 (*t_ox_*). The system of ODEs is given as:

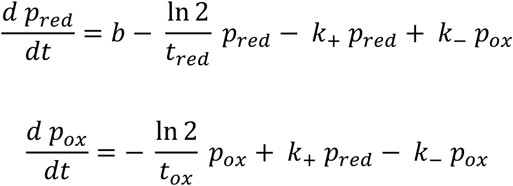

Under equilibrium assumptions the ratio *x* of *p_ox_* to *p_red_* relates to the half-lives according to the following relation:

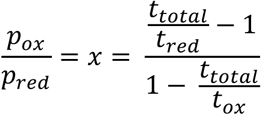

Here, ***t_total_*** refers to the half-life of the total p21 proteins. Figure S6 visualizes this relation for different values of ***t_ox_*** and depending on the ratio ((expressed as the fraction of C41-oxidised 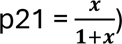. The half-life of reduced p21 was fixed to ***t_red_*** = 5 hours.

### Quantification and Statistical Analysis

Statistical analyses were performed using Prism 10 (GraphPad) with the specific tests indicated in the figure legends. n denotes the number of biological replicates, and N refers to the number of individual measurements. Unless stated otherwise, all experiments were independently repeated at least three times. p-values are indicated in the figures, and values ≤ 0.05 were considered statistically significant. For quantitative Western blot analyses, signal intensities were first normalized to the corresponding loading control and then to the experimental control.

For the BTD data, raw peptide abundances (S/N) were normalized by sample median, log2 transformed and centred at zero per replicate batch. The mean and standard deviation were calculated across all biological and technical replicates. For the total proteome data, the normalized abundances were exported from Proteome Discoverer, log2 transformed, centred at zero per replicate batch and averaged across the two biological replicates. The mean BTD data were then corrected for the mean total protein changes across the time points using linear regression. The protein data were the independent variable (x) whilst the BTD data were the dependent variable (y) per peptide across time points; the residuals of the linear regressions were the BTD changes not attributed to total protein changes. To identify peptides with maximum change at G2/M, S and G1 cell cycle phases, Pearson correlation coefficients and p-values were calculated against defined binary reference profiles in Excel by calculating the t-statistic and using the T.DIST.2T function. Related to Figures 1, 2 and Table S1.

## Supplemental Table Titles

**Table S1 related to Figure 1: Lists BTD, total proteome, and GO analysis data.** Spreadsheet organized in six tabs covering the legend, all MS-data, dynamically oxidized cysteines, GO – G2/M, GO – S, and GO – G1 analysis.

---

**Figure S1 related to Figure 1.**
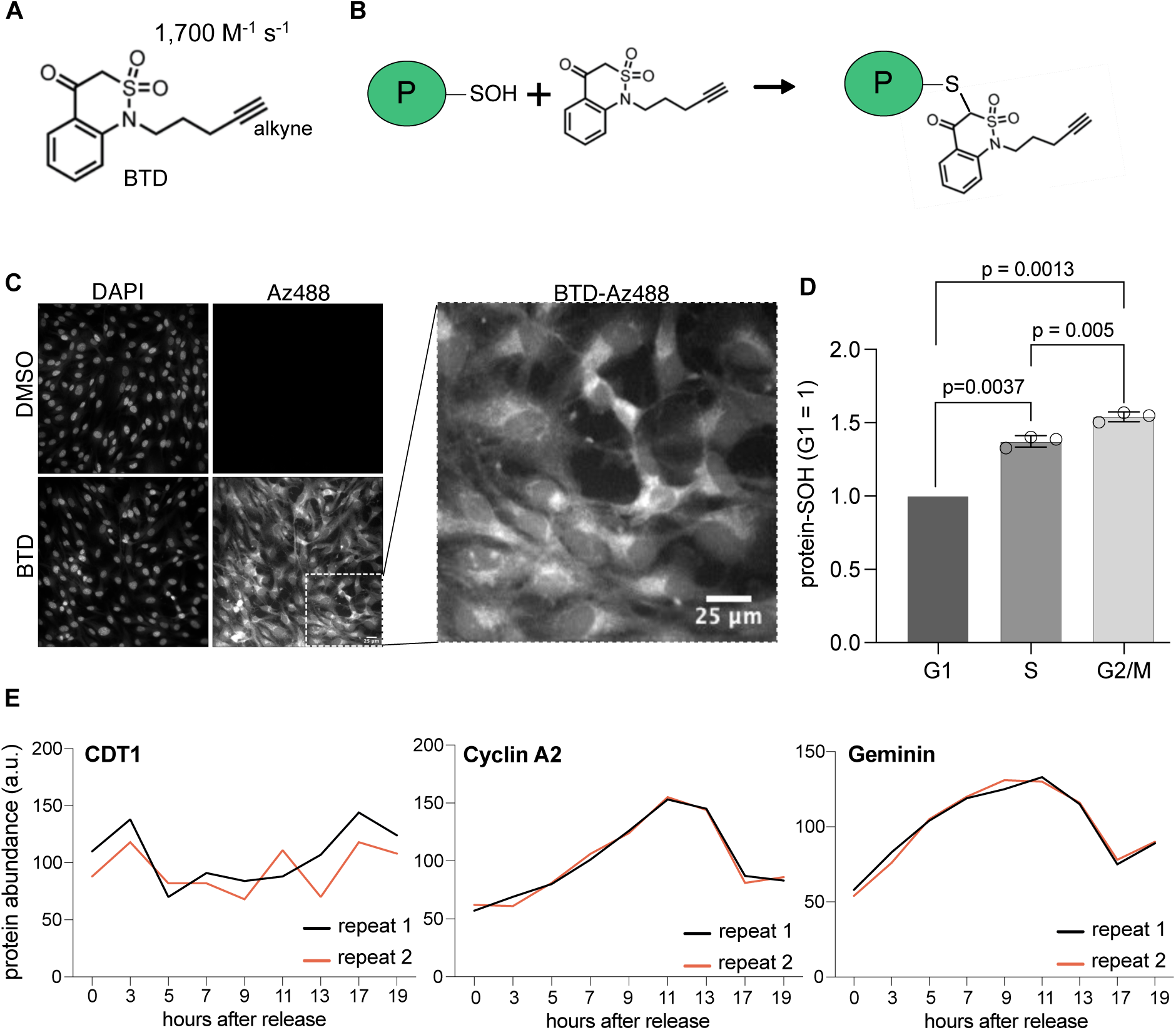
BTD Labeling Detects Increased S-Sulfenylation During Cell Cycle Progression. (**A**) Chemical structure and reaction rate of the sulfenic acid-specific probe BTD, which contains an alkyne group for downstream click-chemistry applications. (**B**) Reaction of BTD with protein sulfenic acids (P-SOH) in living cells. (**C**) *In situ* staining of sulfenic acids in RPE-1 cells treated with BTD or DMSO, using azide dye Az488 clicked to the alkyne moiety of BTD. DNA stained by DAPI. Scale bar, 25 μm; n=3. (**D**) Total S-sulfenylation (protein–SOH), detected by BTD–Az488 and flow cytometry, shown relative to G1 phase. Bars represent mean ± SD; Statistical significance according to one-sample (G1 versus S, G2/M) and paired t-test (S versus G2/M), n=3. (**E**) Relative changes in cell cycle protein abundance determined by quantitative proteomics in RPE-1 cells show high synchronicity between biological replicates (see also Figure 1B). Profiles for all identified proteins are provided in Table S1.

**Figure S2 related to Figure 2:**
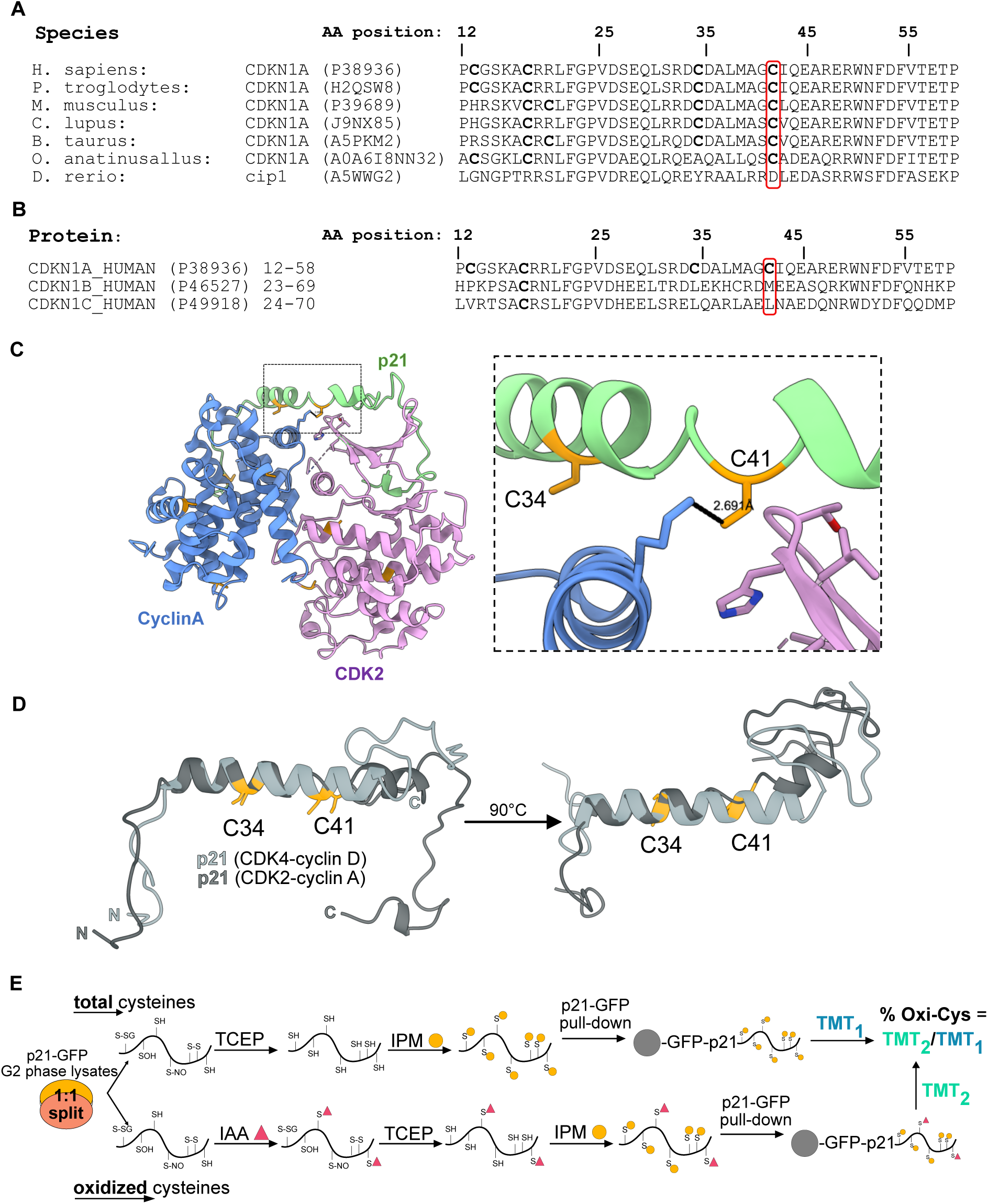
Conservation and Positioning of C41 in complex with CDK2-Cyclin A. (**A**) Alignment of CDKN1A (p21) across vertebrates highlighting the evolutionary conservation of C41, generated using ProViz. Cysteines are highlighted in bold. (**B**) Clustal Omega alignment of CIP/KIP family members in *H. sapiens* showing that C41 is unique to CDKN1A. Cysteines are highlighted in bold. (**C**) Homology model of the p21 (green)-CDK2 (rose)-cyclin A (blue) complex. The inset highlights the position of C41 in p21 relative to cyclin A and CDK2, with inter-residue distances indicated. (**D**) Structural alignment of p21 bound to CDK4–cyclin D (light grey; from Figure 2C) and CDK2–cyclin A (dark grey; from Figure S2C), aligned at the N-terminal end of their α-helices. C34 and C41 are shown in orange. (**E**) Mass spectrometry workflow to determine p21 oxidation stoichiometry. RPE-1 p21-GFP cells synchronized in G2 were split into total cysteine (top) and oxidized cysteine (bottom) samples. For total cysteines, proteins were reduced (TCEP) and alkylated (IPM). For oxidized cysteines, reduced cysteines were first blocked (IAA), then oxidized cysteines were reduced (TCEP) and alkylated (IPM). p21-GFP was enriched using GFP-binder beads, digested, TMT-labelled, and samples were combined. TMT ratios of IPM-labelled peptides indicate cysteine oxidation stoichiometry.

**Figure S3, related to Figure3:**
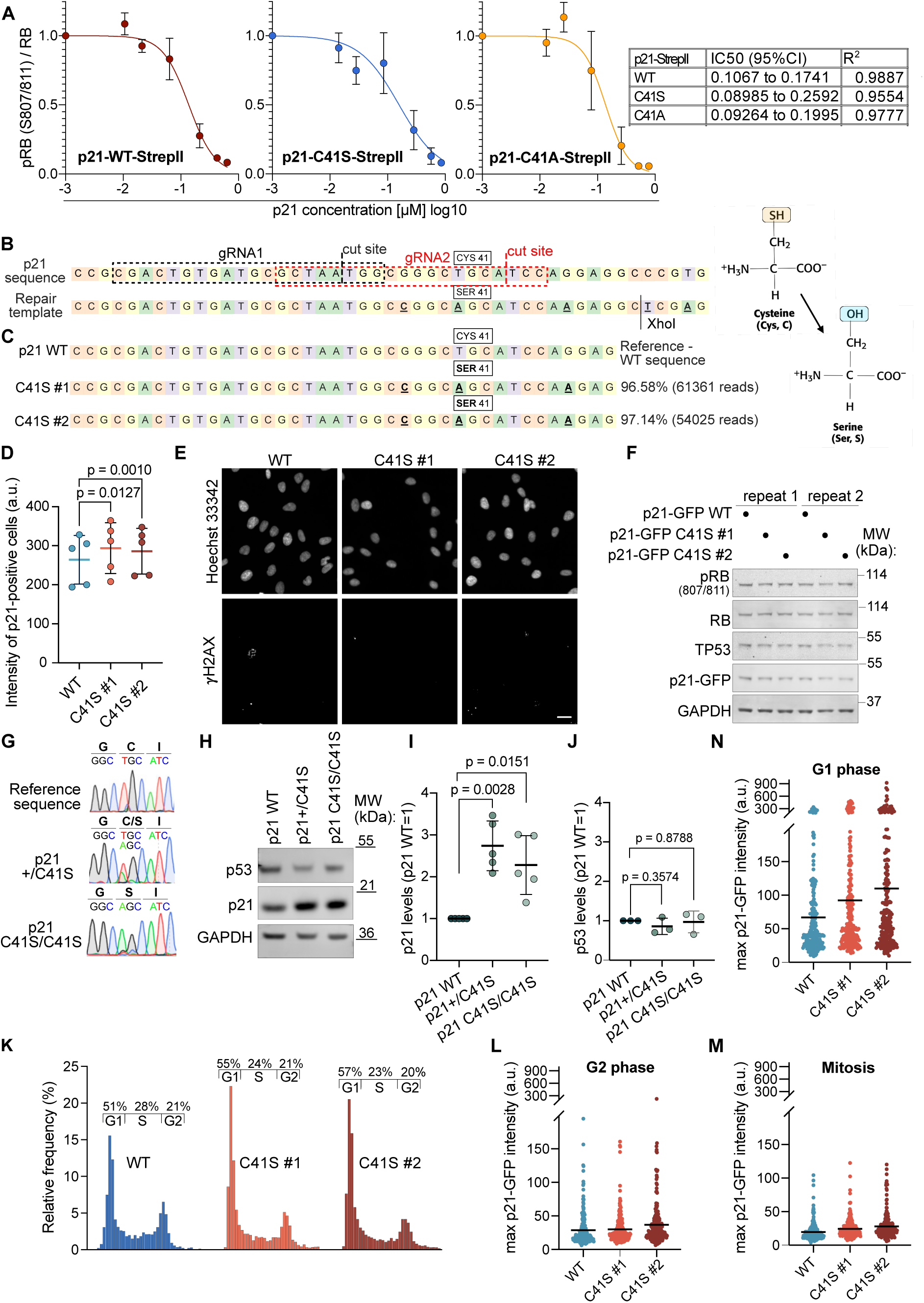
Verification of C41S Mutagenesis and Characterization of p21-GFP WT and C41S Clones. (**A**) *In vitro* kinase assay using recombinant CDK2–cyclin A incubated with increasing concentrations of p21-WT, C41S, or C41A-StrepII, and RB (aa 773–928) as substrate (mean ± SEM; n = 3). IC_50_ values were determined by non-linear regression with variable slopes. (**B**) Top: DNA sequence of p21 showing the guide RNAs used for C41 mutagenesis. Cas9 cut sites are indicated with dashed lines (black for gRNA1, red for gRNA2). Structures of cysteine and serine are shown for reference. Bottom: Repair template used to replace cysteine 41 (CYS41) with serine (SER41), including additional silent mutations (bold, underlined) for genotyping and XhoI digestion. (**C**) DNA sequences of RPE-1 p21-GFP WT and C41S clones determined by amplicon-NGS. Top: p21 WT sequence flanking CYS41. Bottom: Representative alleles and frequencies for C41S #1 and #2 clones. Mutated bases are bold and underlined. (**D**) Quantification p21-GFP intensity from p21-positive RPE-1 cells shown in Figure 3A and 3B (mean ± SD; n = 5). Significance by one-way ANOVA. (**E**) Immunofluorescence of RPE-1 p21-GFP WT and C41S cells stained for γH2AX and DNA (Hoechst 33342). Scale bar, 50 µm. Quantification shown in Figure 3C. (**F**) Western blot of total lysates from asynchronous RPE-1 p21-GFP WT and C41S cells comparing p21-GFP, p53 and RB expression (n=2). (**G**) Sanger sequencing of C41S mutagenesis in MCF10A cells showing conversion of TGC (cysteine) to AGC (serine). Heterozygous clone (p21^+^/C41S) shows both T and A peaks; homozygous clone (C41S/C41S) shows only A. (**H**) Western blot of MCF10A WT, p21^+^/C41S, and C41S/C41S lysates probed for p53, p21, and GAPDH. (**I, J**) Quantification of p21 and p53 protein levels from (H) (mean ± SD; n=5 for p21, n=3 for p53). Significance by one-sample t-tests. (**K**) Frequency distribution of integrated Hoechst 33342 fluorescence intensities from equal numbers of RPE-1 p21-GFP WT and C41S cells to determine cell cycle distribution (n=4). (**L-N**) Maximal p21-GFP intensities in single RPE-1 cells in G2 (L), mitosis (M), and G1 (N) from WT and C41S cells presented in (Figure 3G). Mean values are indicated. (n=2, N=220).

**Figure S4, related to Figure 4:**
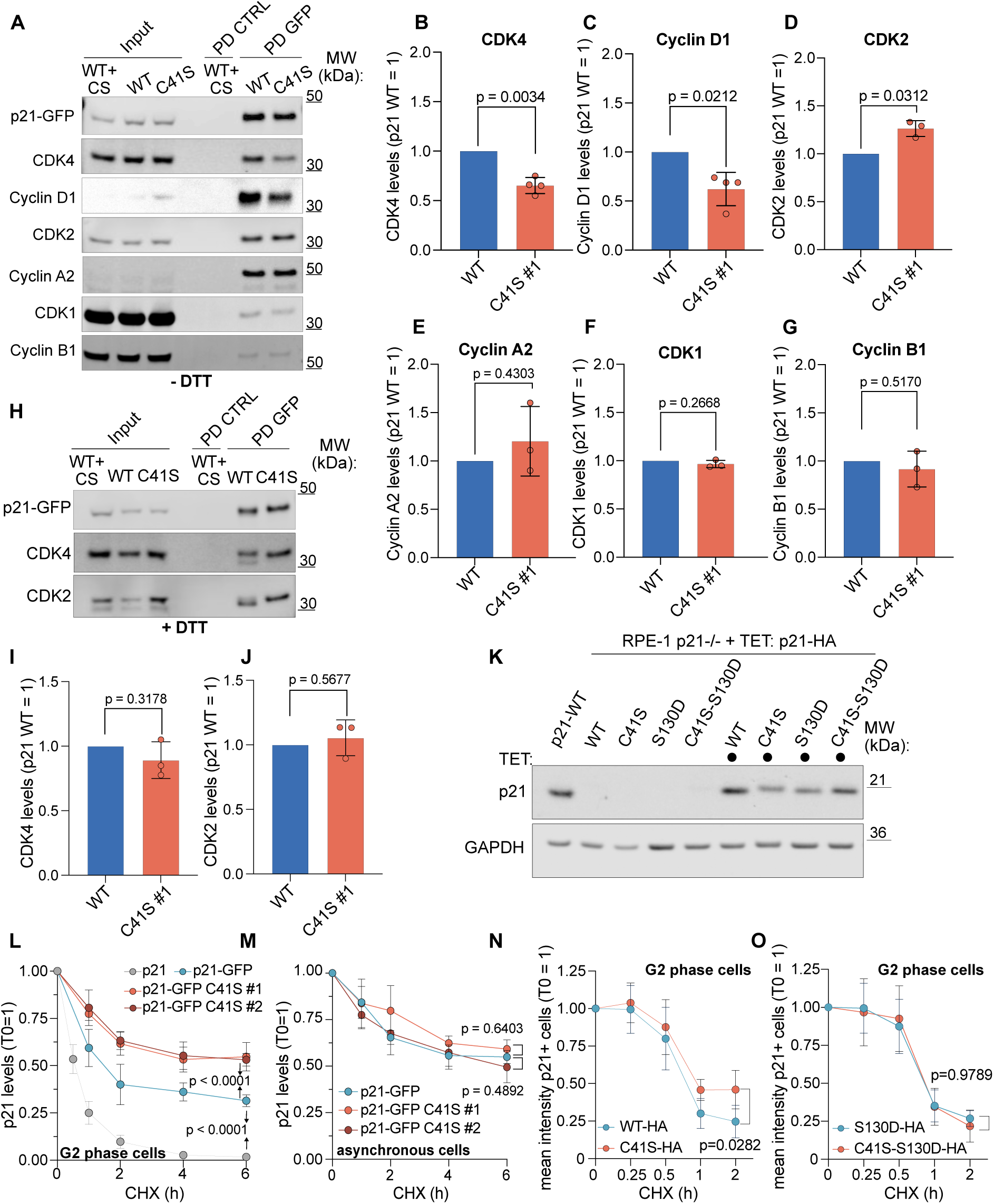
p21 C41S Binding to Cyclin-CDK Complexes and p21 Degradation. (**B-G**) Quantification of data shown in (A) normalized to p21-WT for CDK4 (B), cyclin D1 (C), CDK2 (D), cyclin A2 (E), CDK1 (F), and cyclin B1 (G). Significance by one-sample t-test (mean ± SD; n = 4 for B–C, n = 3 for D–G). (**H**) Western blot as in (A) but with 10mM DTT added to the extract. (**I, J**) Quantification of CDK4 (I) and CDK2 (J) binding from (H), normalized to p21-WT. Significance by one-sample t-test. Note: under reducing conditions, no significant dioerence is observed between p21-WT and C41S (mean ± SD; n = 3). (**K**) Western blot of total lysates from asynchronous RPE-1 p21^-^/^-^ cells rescued with tetracycline-induced p21-WT-HA, p21-C41S-HA, p21-S130D-HA, and p21-C41S-S130D-HA, all expressed at levels comparable to endogenous p21. GAPDH serves as loading control. (**L**) Decay curves of p21 during G2 from Western blot data as in Figures 4G and 4H. Significance at 6h by two-way ANOVA (mean ± SD; n = 3, N = 4). (**M**) Decay curves of p21 in asynchronous cells as from G2 cells shown in Figure 4H. Significance at 6 h by two-way ANOVA (mean ± SD; n = 3, N = 4). (**N, O)** Decay curves in G2 cells of HA-tagged p21-WT and p21-C41S (N) or p21 WT + S130D and p21 C41S + S130D mutants (O), measured by anti-HA immunofluorescence and Hoechst 33342–based DNA content gating. Significance at 2 h by two-way ANOVA (mean ± SD; n = 6).

**Figure S5 related to Figure 5:**
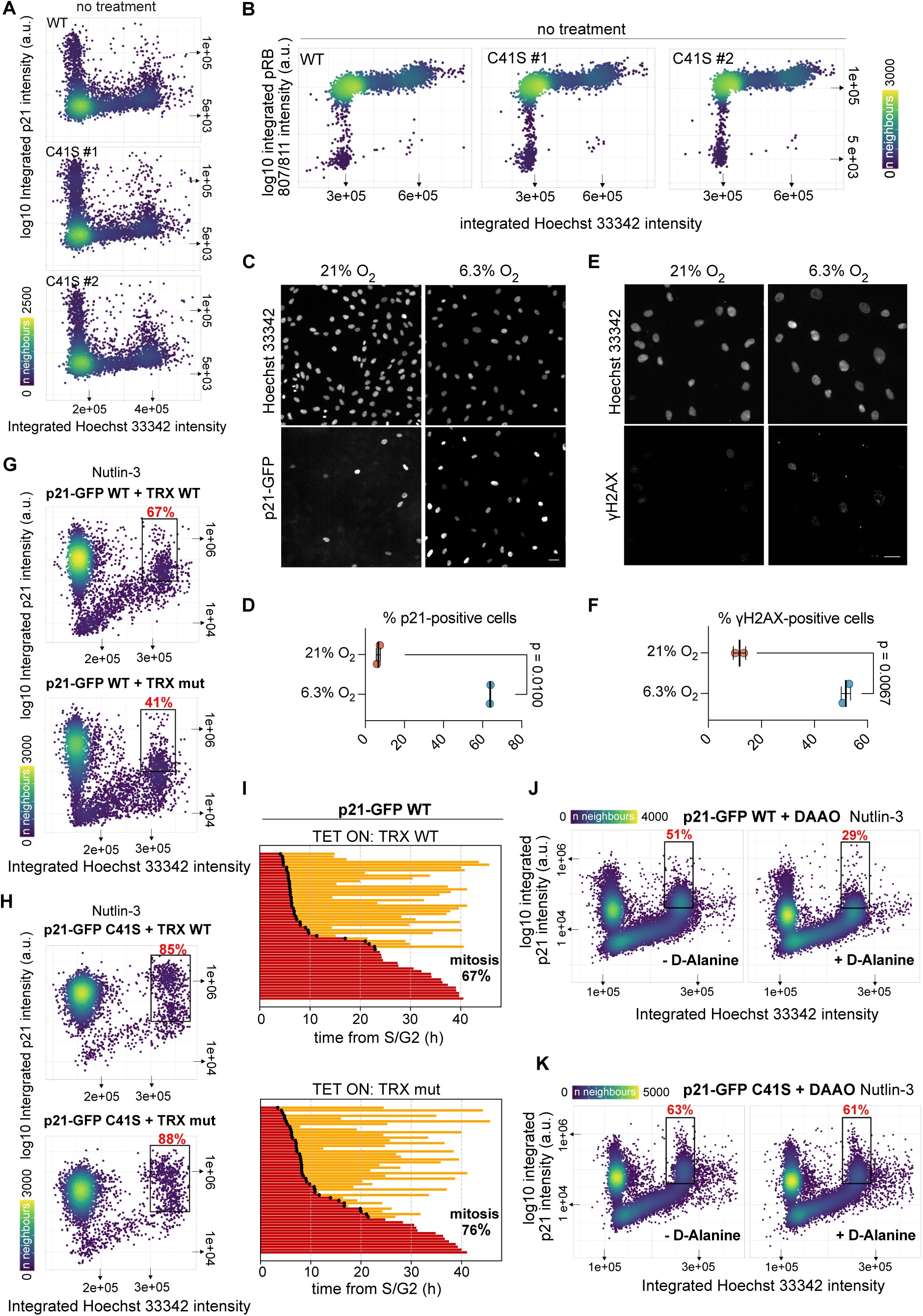
C41 Redox State Modulates p21 Stability and Sensitivity to Cellular Stress. (**A**) Bivariate density plots showing DNA content and p21 levels in WT and C41S cells under unperturbed conditions. Related to Figure 5A. (**B**) Bivariate density plots showing DNA content and pRb S807/811 staining in WT and C41S cells under unperturbed conditions. Related to Figure 5D. (**C**) Fluorescence microscopy of p21-GFP and DNA (Hoechst 33342) in RPE-1 cells cultured at 21% O_2_ (hyperoxia) or 6.3% O_2_ (normoxia). Scale bar, 50 µm. (**D**) Quantification of (C) showing the increased percentage of p21-positive cells at 6.3% O_2_. Statistical significance by paired t-test (mean ± SD; n = 2). (**E**) Immunofluorescence of p21-GFP WT cells stained for γH2AX and DNA (Hoechst 33342) after culture at 21% or 6.3% O_2_. Scale bar, 50 µm. (**F**) Quantification of γH2AX-positive cells from (E). Statistical significance by paired t-test (mean ± SD; n = 2). (**G, H**) Bivariate density plots of DNA content and p21-GFP intensity in p21-GFP WT (G) and C41S (H) cells expressing HA-VHH4-TRX WT or HA-VHH4-TRX mut after 24 h of treatment with 1 μM Nutlin-3. Gating and percentage of p21^high^ cells in G2 are indicated. Corresponding quantification is presented in Figures 5I and 5J. (**I)** Single-cell tracks from live-cell imaging of RPE-1 p21-GFP WT cells expressing equal levels of TRX WT or TRX mut (as in Figure 5H). TRX expression was induced during S phase together with 1 μM Nutlin-3 to increase p21 levels. Tracks show progression through G2 (red), mitosis (black dot), and entry into the next cell cycle (yellow). Only tracks lasting at least 5x the average duration of unperturbed G2 (4.78 h, see Figure 3G) were included to reduce tracking bias. Percentage of cells entering mitosis is indicated. (**J, K**) Bivariate density plots of DNA content and p21-GFP intensity in p21-GFP WT (J) and C41S (K) cells expressing HA-VHH4-DAAO, after 3 h of treatment with 5 μM Nutlin-3 with or without D-alanine. Gating and percentage of p21^high^ cells in G2 are indicated. Corresponding quantification is shown in Figures 5M and 5N.

**Figure S6 related to Figure 7:**
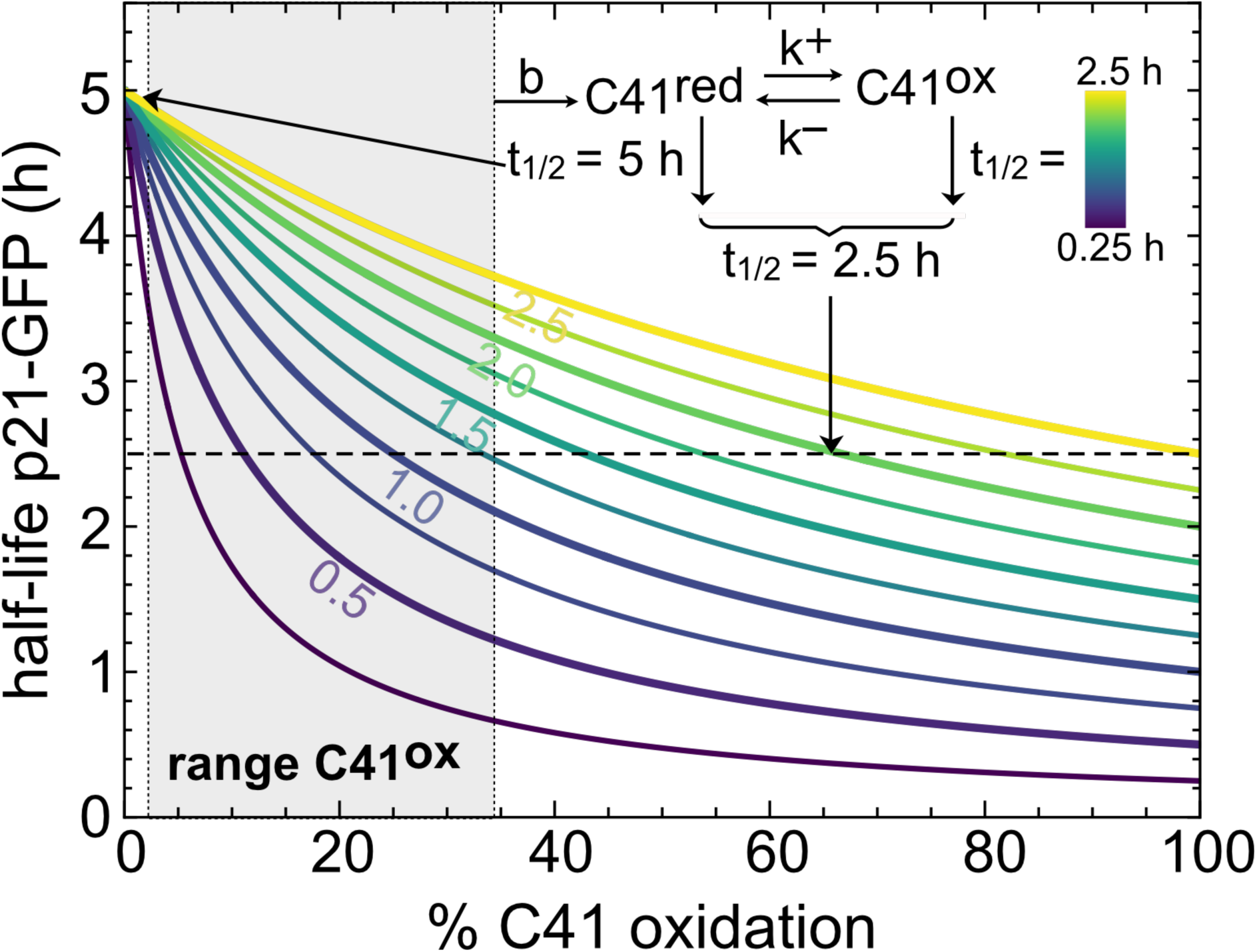
Eaect of C41 Oxidation on p21 Half-Life. Ordinary Dioerential Equation (ODE) model describing the dynamic equilibrium between C41-reduced and C41-oxidized p21 to evaluate how varying degradation rates, expressed as half-life (t), influence overall p21 stability. The half-life of reduced p21 (C41^red^) was fixed at 5 hours, based on the experimentally determined half-life of non-oxidizable p21-GFP C41S (Figure 4). The shaded grey area denotes the range of experimentally determined C41 oxidation stoichiometry by mass spectrometry (Figure 2F). The experimentally observed half-life of total p21-GFP (∼2.5 h; Figure 4I) is indicated by a dashed horizontal line. At the average oxidation level of ∼12%, the model projects a half-life of ∼0.5 h for p21-GFP fully oxidized on C41 (C41^ox^), suggesting that the total p21 pool can be rapidly turned over due to continuous oxidation of reduced p21 during G2 phase.

**Table S2 related to STAR Methods:**
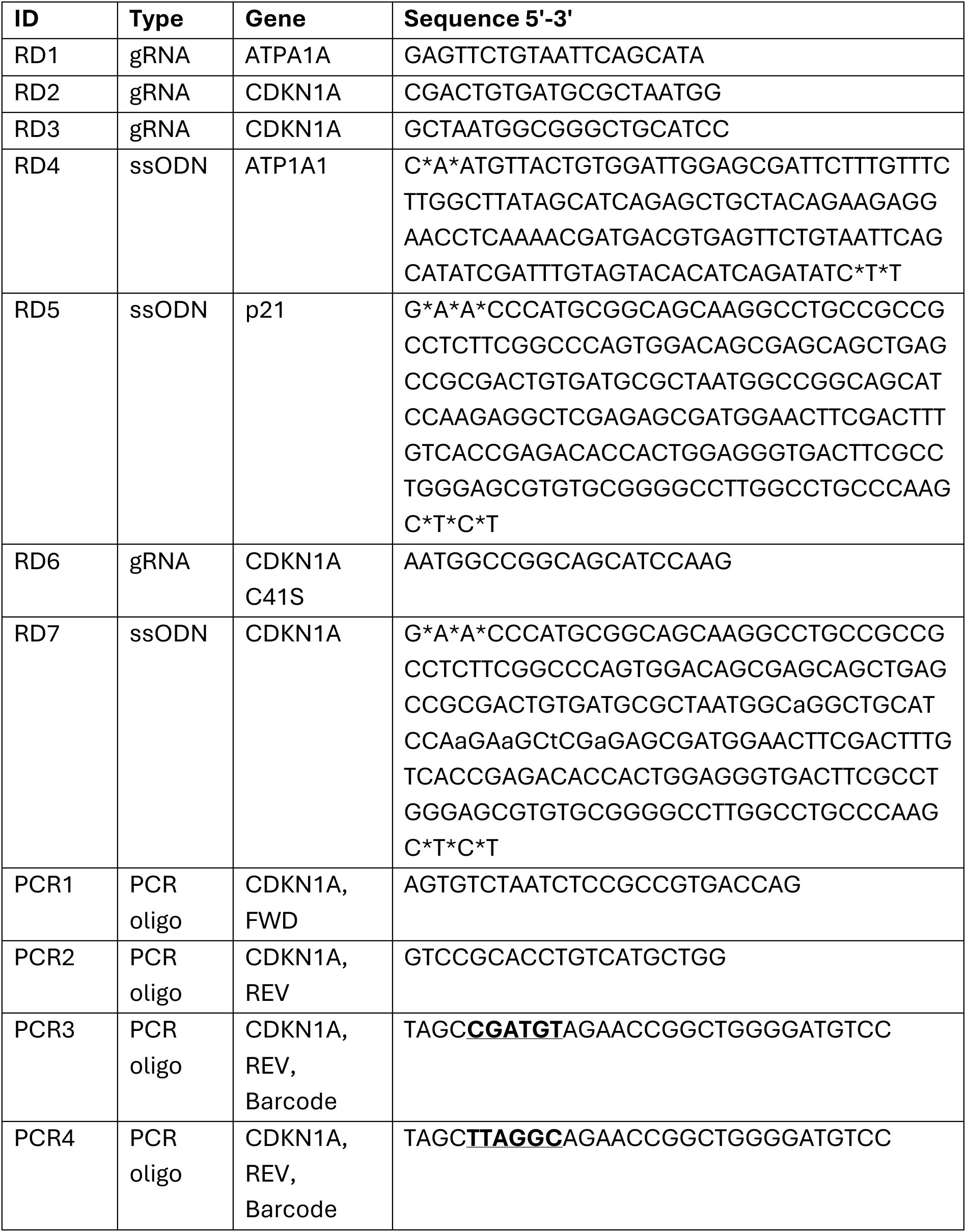
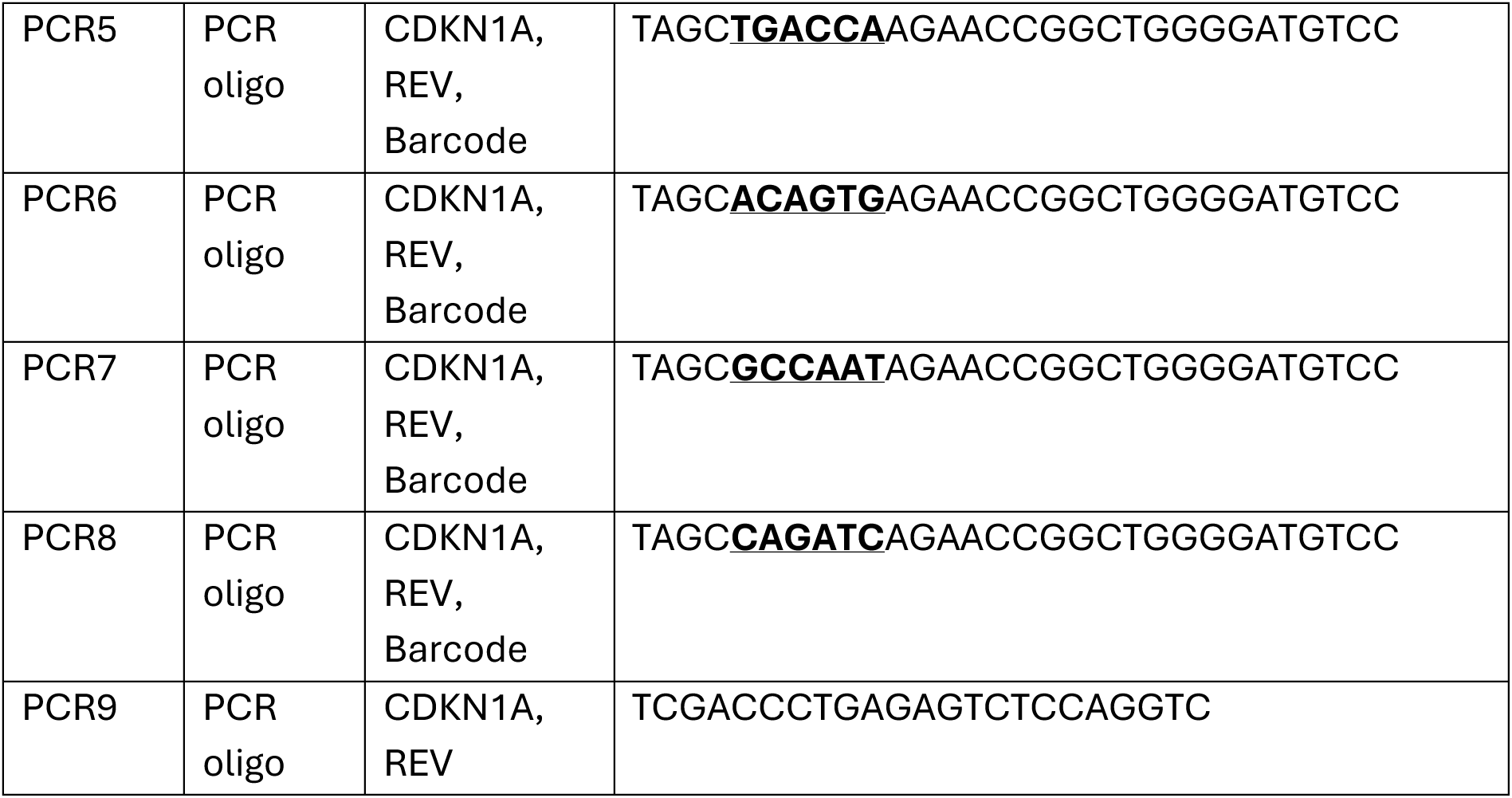
Lists DNA oligonucleotides used in this study. Guide RNAs (RD), repair templates (ssODN), and PCR primer sequences (PCR) used in this study. * denotes phosphorothioated nucleotides. FWD = forward, REV = reverse, barcode underlined and in bold.

## Notes

### Competing Interest Statement

The authors have declared no competing interest.

### Summary of Updates

This is the last author submitted version before acceptance. Updates were made to most figures and the text. This includes changes to the title, abstract, and the addition of two authors.

